# In situ crystalline structure of the human eosinophil major basic protein-1

**DOI:** 10.1101/2024.10.09.617336

**Authors:** Jie E. Yang, Joshua M. Mitchell, Craig A. Bingman, Deane F. Mosher, Elizabeth R. Wright

**Author notes:** Declaration of interest: none.

## Abstract

Eosinophils are white blood cells that participate in innate immune responses and have an essential role in the pathogenesis of inflammatory and neoplastic disorders. Upon activation, eosinophils release cytotoxic proteins such as major basic protein-1 (MBP-1) from cytoplasmic secretory granules (SGr) wherein MBP-1 is stored as nanocrystals. How the MBP-1 nanocrystalline core is formed, stabilized, and subsequently mobilized remains unknown. Here, we report the *in-situ* structure of crystalline MBP-1 within SGrs of human eosinophils. The structure reveals a mechanism for intragranular crystal packing and stabilization of MBP-1 via a structurally conserved loop region that is associated with calcium-dependent carbohydrate binding in other C-type lectin (CTL) proteins. Single-cell and single-SGr profiling correlating real-space three-dimensional information from cellular montage cryo-electron tomography (cryo-ET) and microcrystal electron diffraction (MicroED) data obtained from non-activated and IL33-activated eosinophils revealed activation-dependent crystal expansion and extrusion of expanded crystals from SGr. These results suggest that MBP-1 crystals play a dynamic role in the release of SGr contents. Collectively, this research demonstrates the importance of *in-situ* macromolecular structure determination.

## Main

Eosinophils are specialized immune cells derived from hematopoietic stem cells that combat infections, participate in the pathogenesis of allergic inflammatory diseases such as asthma, and maintain immune homeostasis^1^. Mature eosinophils are readily identified by the presence of large cytoplasmic secretory granules (SGr) that appear reddish-orange when stained with eosin dye^2^. SGr store a number of mediators^3^, including highly abundant cationic proteins that dominate the human eosinophil proteome^4^. SGr in mature eosinophils contain a central electron-dense nanocrystal core surrounded by a more translucent matrix material^5^. Eosinophil major basic protein-1 (MBP-1), a highly positively charged protein (117-residues with a calculated pI of 11.4, and net charge of 15 at pH 7)^6^, comprises the crystalline core^7^. MBP-1 is derived from the proform, proMBP-1 (222-residues), also known as bone marrow proteoglycan-2 (PRG2)^8^, and matures through cleavage of the propiece. Soluble MBP-1 is cytotoxic and causes membrane disruption to pathogens and vertebrate cells^9^. Both proMBP-1 and the isolated nanocrystalline cores of MBP-1 are nontoxic^8,10^. It has been proposed that eosinophils, the secretory apparatus, and SGrs are protected from non-specific cytotoxic damage by retaining proMBP-1 until just prior to its cleavage to MBP-1, and then MBP-1 compaction and crystallization into nanocrystals^10^. In other words, MBP-1 crystals are stored safely in eosinophil SGrs.

A key event in the pathology of eosinophil-mediated inflammatory diseases is the process of degranulation. Following eosinophil activation, SGrs mobilize and release their granule content into the extracellular space^11^. Four mechanisms of degranulation have been identified^11^, including piecemeal degranulation^12^, cytolysis^13^, compound exocytosis, and classical exocytosis^14^. Eosinophils may employ multiple degranulation pathways simultaneously and preferentially based on the activating agent^11^. Piecemeal degranulation is a progressive and selective secretion process of macromolecules from the SGr coordinated by structures known as eosinophil sombrero vesicles (EoSVs)^3^. EoSVs interact with and bud from SGr, transporting granule-derived products including MBP-1 to the extracellular space^15^. Nanocrystals dissolve at pHs below 4^7^, and SGrs undergo acidification during degranulation^16^, suggesting a mechanism whereby MBP-1 is mobilized for transport by EoSVs. Classical exocytosis proceeds with the release of the entire granule through fusion of the granule with the cell’s plasma membrane. Compound exocytosis is similar to classical exocytosis, but involves an extra step of granule-to-granule fusion prior to extracellular release^14^. Lastly, cytolysis^13^ occurs when the activated eosinophil undergoes rapid nonapoptotic cell death to release intact SGr via rupturing of the plasma membrane. Little is known about the mechanism of SGr nanocrystal formation and its subsequent mobilization during degranulation. This is partly due to the elusive nature of the MBP-1 intragranular nanocrystal structure. Prior protein crystallography of acid-solubilized and recrystallized eosinophil MBP-1^17,18^ demonstrated an unusual C-type lectin (CTL) domain fold that lacked a site to bind carbohydrates in a calcium-dependent manner. Probing isolated SGr by X-ray-free-electron laser crystallography (XFEL)^10^ revealed yet another crystal lattice distinct from the *in vitro* structure, indicating that intragranular MBP-1 crystals may be structurally different.

To explore SGr degranulation, we cryogenically preserved non-activated and activated eosinophils in their native states. We directly probed the structure of eosinophil intragranular nanocrystals in the unperturbed cellular environment at the single granule level using correlative cryo-focused ion beam (cryo-FIB) milling ^19^, microcrystal-electron diffraction (MicroED)^20^, and montage cryo-electron tomography (cryo-ET)^21^. We report the structure of MBP-1 in its native nanocrystalline form and show that the configuration of calcium-dependent carbohydrate binding site present in other proteins with CTL domains is also conserved by MBP-1, yet functions to accommodate intragranular crystalline packing. The conformation of MBP-1 in the native nanocrystalline form is distinct from what was determined in previous *in vitro* studies. Correlative single-cell and single-SGr profiling revealed a mechanism for activation-dependent intragranular crystal expansion and reaffirms that there is a diversity of degranulation pathways. These results suggest that MBP-1 crystals play a dynamic role in SGr content release.

### Native human eosinophil crystalline core and structure determination

To directly study the structure of secretory granules (SGr) in the native environment of human eosinophils, we used eosinophils purified from donors’ blood and deposited them onto fibrinogen-coated TEM grids. We used interleukin-33 (IL33) as a cytokine activator; IL33 is known as a key cytokine for innate-type mucosal immunity^22^, and stimulates eosinophil adhesion, degranulation, and chemotaxis^23^. IL33 causes eosinophils to adhere via ITGAM/ITGB2 integrin to fibrinogen-coated glass coverslips or TEM grids with a highly flattened morphology^24^ that supports electron beam penetration for cryo-EM imaging and crystal diffraction. Non-activated, resting eosinophils were captured on fibrinogen-coated TEM grids or coverslips. Live-cell imaging prior to plunge freezing and initial low-dose cryo-EM (Extended Data Fig. 1, 2) demonstrated little damage or unwanted stimulation to non-activated eosinophils (Extended Data Fig. 1a-b, 1d, 2a-c) and consistent activation of IL33-activated eosinophils (Extended Data Fig. 1c, 1e-g, 3a-b). Non-activated eosinophils displayed a typical sphere-like cell shape (Extended Data Fig. 1a-b, 2a-b), and most of their SGr were in regions too thick for electron beam penetration, resulting in poor crystal diffraction (Extended Data Fig. 2c-d). In contrast, IL33-activated eosinophils were flattened and SGr were TEM accessible in thin extensions of the cell periphery (Extended Data Fig. 1c, 1e-g, 3b-c). However, even in flattened cell edges, intragranular crystalline cores showed weak diffraction (Extended Data Fig. 3c, white arrowhead).

Unexpectedly, membrane-less free crystalline cores (Extended Data Fig. 3-6, asterisks and white arrows) were observed in IL33-activated cells. These were dispersed in the cell edge extensions protruding away from the main body (Extended Data Fig.1c, 1e-g, red arrows, 3b-c, 4a-b). In these regions, the plasma membrane boundary was very thin and barely discernable under TEM (Extended Data Fig. 3b, 4a-b, asterisks). The presence of membrane-less crystals could be a consequence of eosinophil activation or an artifact of grid-deposition-freezing steps^25,26^. We therefore examined non-activated eosinophils on carbon-coated grids via low dose TEM. Except for a small number of cells (roughly 5-10%) that exhibited activated cell flattened morphology, free crystals were absent in non-activated eosinophils (Extended Data Fig. 2). Eosinophil activation by handling has been observed previously^27^. Therefore, we concluded that the free crystals resulted from either handling of the cells, IL33-stimulation, or a combination of the two. To differentiate intra- or extra-cellular locations of free crystals after IL33 activation, we conducted correlative cryo-light and electron microscopy (cryo-CLEM)^28^. A low toxicity far-red carbocyanine dye^29^ was used to stain the cytoplasmic membrane (Extended Data Fig. 6) and reached an equilibrium binding affinity between cytoplasmic and intracellular membrane structures including the delimiting membrane of SGr. Autofluorescence of the SGr matrix (green) from granule-associated flavins^30^ (Extended Data Fig. 6c-e) produced a colocalized signal with that of the SGr membrane (red). Consistently, membrane-less naked crystals were apparent, observed along or near the flattened membrane edges of two-thirds of the activated eosinophils imaged (119 out of 190), indicative of active release from the cell (Extended Data Fig. 6f, g-i). Activated eosinophils are well-known to undergo cytolytic degranulation to release SGr with intact membranes into the extracellular space concomitant with disintegration of the plasma membrane^11,13^. Our results suggest that naked crystals can also be released from tightly adherent IL33-activated eosinophils (Extended Data Fig. 5c-f, white arrows, 6f). We cannot exclude the possibility that the free crystals arise because of increased susceptibility of SGr membranes to loss of integrity associated with damage during the blotting/plunge-freezing step^25,26^. Crystals in membrane compartments are known to make membranes susceptible to lysis^31^. Importantly, despite the lack of delimiting SGr membranes (Extended Fig. 3c-f), the free crystals displayed lattice packing (Extended Data Fig. 3d-e) and diffracted up to 3 Å (Extended Data Fig. 3f). As shown in Extended Data Fig. 3g, the diffraction pattern quickly faded after the application of small electron doses (∼0.25 e/Å^2^), indicating that either crystal disassembly was already underway or they had a less perfect crystal lattice packing^32^. Thus, these observations of the naked crystals provided a strong rationale for determining the structure of the intragranular crystalline core in unperturbed eosinophils.

### *In-situ* structure of human eosinophil granule major basic protein-1 (gMBP-1)

We developed a versatile workflow for *in-situ* investigations of the human intragranular eosinophil major basic protein-1 (gMBP-1) and its crystal packing (Extended Data Fig. 7). Cryo-focused ion beam milling (cryo-FIB) was used to generate 200-250 nm thin lamella per eosinophil (Extended Data Fig. 8) in a non-activated (resting), or IL33-activated state. The cryo-FIB milling exposed granules present in their native cellular context and made them accessible for subsequent cryo-EM structural analysis. Due to their small size, the intragranular nanocrystals were particularly suited for microcrystal electron diffraction (MicroED)^20^. The condensed and crowded SGrs in the cytoplasm required the use of montage cryo-electron tomography (cryo-ET) via montage parallel array cryo-tomography (MPACT)^21^ to map SGr and nanocrystal locations within each eosinophil and associated sub-cellular compartments. We acquired and catalogued intracellular MicroED and montage cryo-ET data for each target to build comprehensive correlative 3D *in-situ* profiles of individual SGr (Extended Data Fig. 7). As shown in Figure 1, SGrs (highlighted in yellow in Extended Data Fig. 8c-f, Fig. 1a-d) provided sufficient contrast for the analysis of intragranular contents by cryo-FIB-SEM (Extended Data Fig. 8a-c), low-dose cryo-EM (Extended Data Fig. 8d-f), MicroED (Fig. 1e), and montage cryo-ET to correlate 3D volumes (Fig. 1b) with associated diffraction data (Fig. 1). We termed this workflow “*in-situ* single granule profiling”. Both electron diffraction (Fig. 1c) and real space analysis via Fast Fourier Transform (FFT) of cryo-FIB milled granules (Extended Data Fig. 8e-f) showed diffraction signals discernable up to 2.9 Å (Fig. 1c).

**Figure 1.**
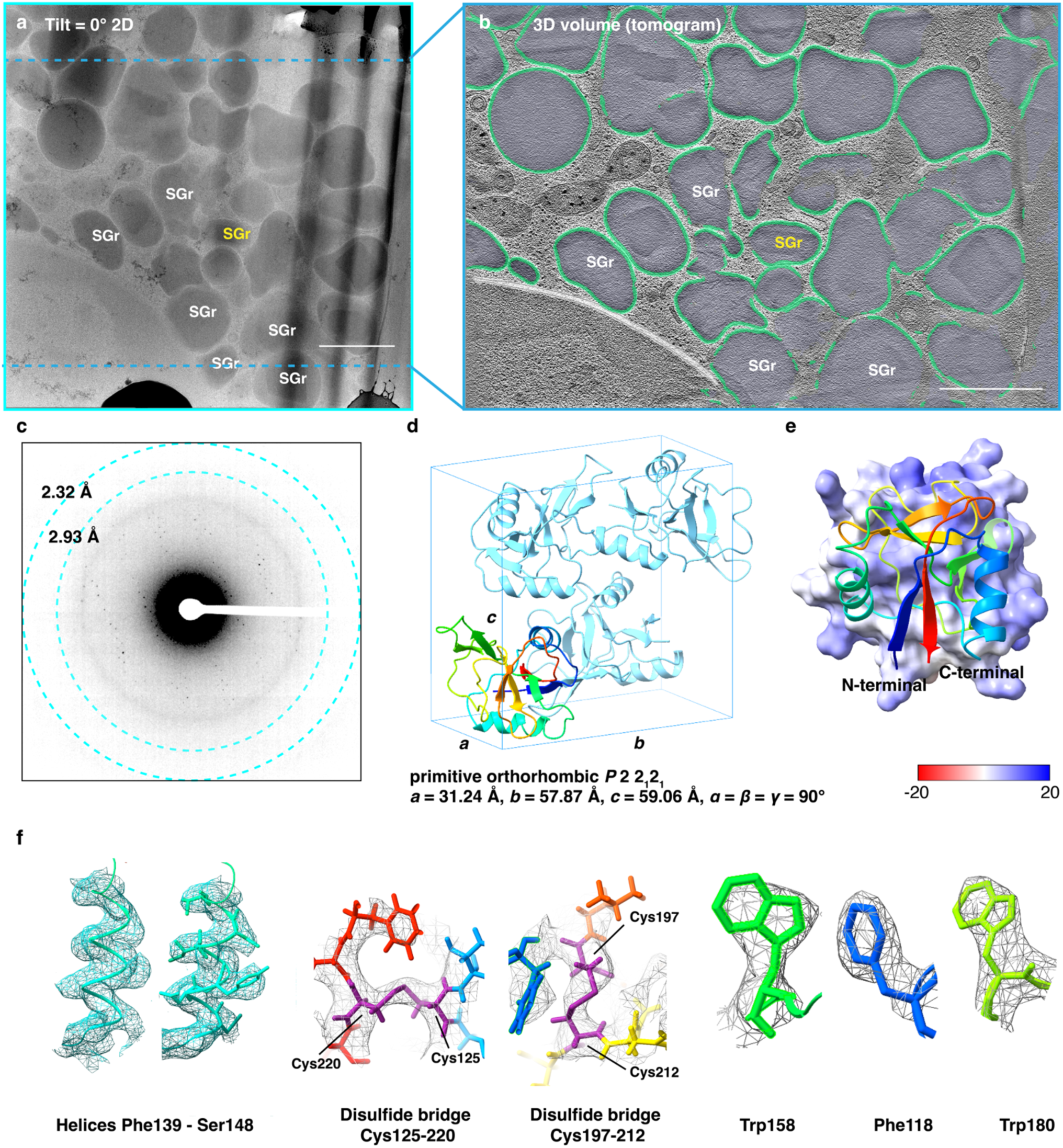
*In-situ* MicroED structure of human eosinophil major basic protein-1 (gMBP-1). **a**, A 2D view of a stitched 3×3 MPACT montage tilt series at the tilt angle of 0° (stage tilt of 9°) of a post-milled eosinophil cell (lamella thickness of 220 nm). **b**, A tomographic slice (2-slice average, thickness of ∼38 nm) of the final reconstructed MPACT tomogram from the tilt series (**a**) with segmented SGr where the delimiting membrane (green) and matrix content (purple) are delineated. The final field of view of the reconstructed tomogram (**b**) is determined by the final high tilt in the tilt series (blue dashed lines in **a**). **c**, A representative diffraction (summed over 5° wedge) from the MicroED dataset of the same highlighted granule (yellow SGr in **a-d**). **d**, The space group of gMBP-1 (PDB code: 9DKZ) in ribbon model representation. **e,** gMBP-1 in a ribbon view (rainbow) with the calculated electrostatic molecular surface (red-white-blue palette, the minimum and maximum potentials of gMBP-1 are -2.87 and 17.53, respectively). **f**, Individual amino acid residue models were overlayed with the 2mF_o_- DF_c_ map (contoured at 1.5 *σ*), coded as in **e**. Scale bars of 1µm in **a-b**.

There were no observable adverse effects to the electron diffraction patterns after a total dose of 2 e/Å^2^, indicative of good preservation of nanocrystals post FIB-milling^33^ (Extended Data Fig. 9). Following standards used to determine optimal dose for MicroED^34,35^, we monitored the fading of diffraction spot intensities. Diffraction spots with intensities that could be reliably indexed persisted up to an accumulated dose of 6.5 e/Å^2^, although some high-resolution diffraction spots (diffraction beyond 4 Å) did start to broaden (Extended Data Fig. 9a). Thus, we optimized the data acquisition scheme to keep the total dose below 6.5 e/Å^2^ (Extended Data Fig. 9b).

MicroED acquisition and analysis were performed on electron-dense SGr or granule-like vesicles on thin lamellae. The small size of cytosolic SGr and their proximity to one another made it difficult to avoid illuminating adjacent crystals at high tilts. As a result, approximately 6 % of the raw datasets (51 out of 829) from non-activated (n = 20 SGrs) or IL33 activated (n = 31 SGrs) eosinophils were of sufficient quality to be indexed. Six nanocrystal datasets of SGr from non-activated eosinophils were merged to reach an overall completeness of 96% (Extended Data Fig. 10a, Table 1) and allowed us to solve the structure of native granule major basic protein-1 (gMBP-1) at a resolution of 3.2 Å (Fig. 1d-f, Extended Data Fig. 10). The unit cell parameters were determined to be *a* = 31.24 Å, *b* = 57.87 Å, *c* = 59.06 Å, *α* = *β* = *γ* = 90°, imposing a primitive orthorhombic symmetry of *P* 2 2_1_ 2_1_, one MBP-1 molecule per asymmetric unit (Fig. 1c). Notably, the intragranular nanocrystals situated inside the cell displayed a distinctly different crystal symmetry than the lattice packing adopted by human MBP-1 purified and re-crystalized *in vitro* (pMBP-1, PDB code: 1h8u)^18^ (space group *C* 2, *a* = 74.33 Å, *b* = 57.49 Å, *c* = 60.96 Å, *α* = *γ* = 90°, *β* = 113.2°) or nanocrystals in isolated SGr probed by X-ray-free electron laser (XFEL)^10^ (reported space group *P* 2, *a* = 53.9 Å, *b* = 25.8 Å, *c* = 59.2 Å, *α* = *γ* = 90°, *β* = 90.2°). To be sure that the intracellular crystals were best described by *P* 2 2_1_ 2_1_, we explored lower symmetry alternatives by rescaling and reintegrating in monoclinic lattice. There was a consistent preference for primitive orthorhombic symmetry. Of note, the unit cell dimensions from the XFEL data had very similar cell constants as gMBP-1, with angle *β* being 90.2°. The observed differences present in the intragranular lattice packing may arise from changes in the granular micro-environment during isolation and experimental processing. As pointed out by the authors^10^, the purified SGrs might have suffered from dehydration.

**Table 1.**
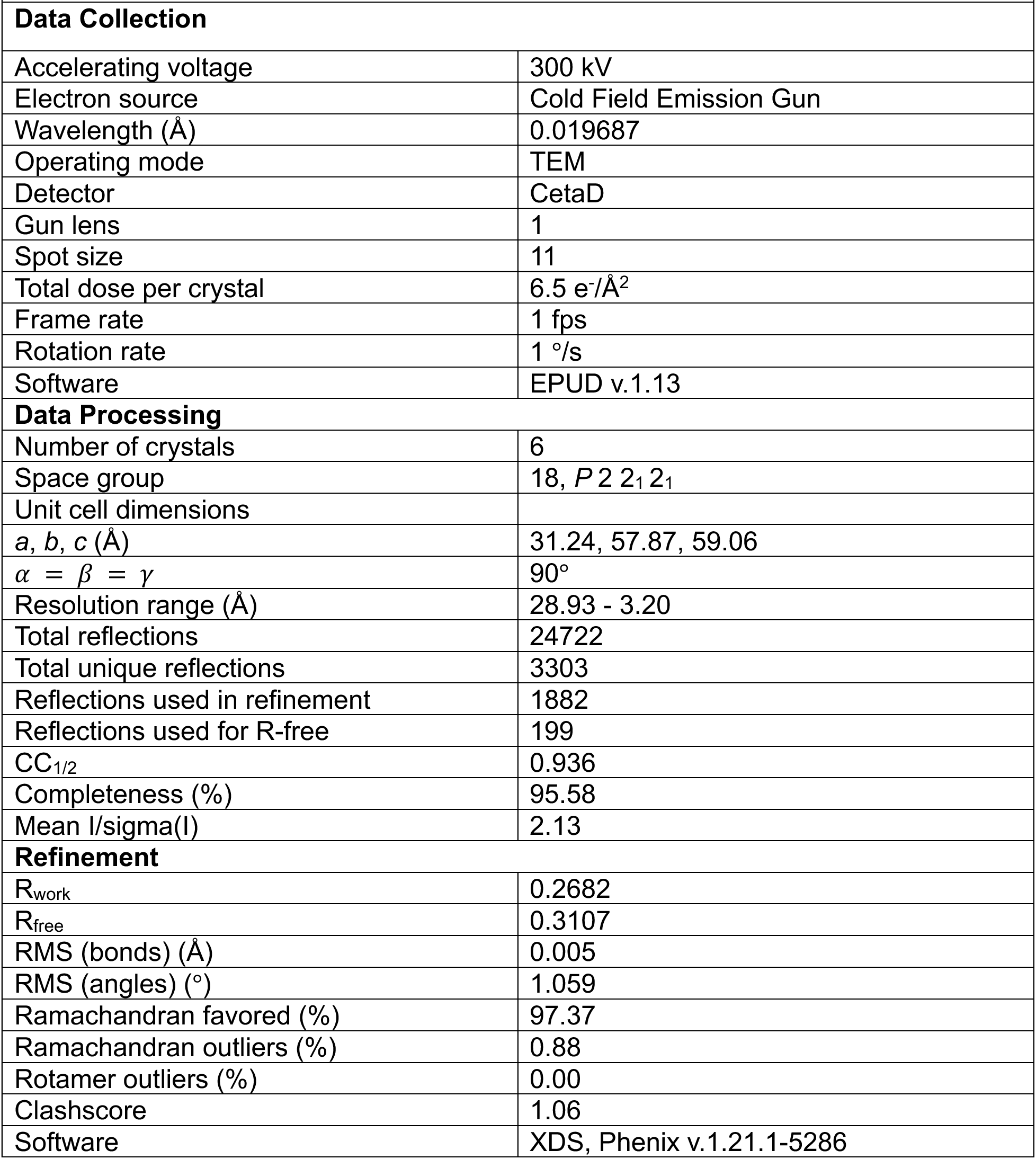
MicroED Data collection and refinement statistics.

The structure of granule MBP-1 (gMBP-1) was determined by molecular replacement (MR)^36^. The phase could be resolved using either the purified and recrystallized MBP-1 monomer (pMBP-1, PDB code: 1h8u.pdb1 chain A)^18^ or an Alphafold predicted model based on the pMBP-1 sequence (1h8u.fa)^18^ (Extended Data Fig. 10b). Similar to pMBP-1^18^, the flexible loop of residues 150-155 (residues numbered starting with the initial methionine of PRG2) was less well defined. The long loop (residues 162-171) in gMBP-1 was poorly resolved, indicating flexibility and less ordered packing in this region. We chose to include it in the final model (PDB code: 9DKZ), although we were less certain about the structure of those residues.

Elsewhere, the map showed well-defined and overall well-resolved density (Fig. 1f). The final structure had crystallographic R-factor (Rwork) and cross-validated R-factor (Rfree) values of 0.26 and 0.31, respectively (Table 1). The overall real space correlation was 0.748, mostly caused by the poor density in the loop regions (residues 150-155, 162-171). As expected^17,18^, the overall surface potential of *in-situ* MBP-1 protein was a highly positive, as expected for a very basic protein (Fig. 2b). The granule MBP-1 (gMBP-1) structure has the architecture of C-type lectin family/domain (CTL), comprised of two *α*-helices and seven *β* strands that form three anti-parallel *β* sheets. Two disulfide bonds (Cys125-220 and Cys197-212, Fig. 1f, Extended Data Fig. 10b) help maintain local conformations by connecting *α*1 and *β*7, and stabilizing *β*6 - Loop6 - *β*7, respectively (Extended Data Fig. 10b).

**Figure 2.**
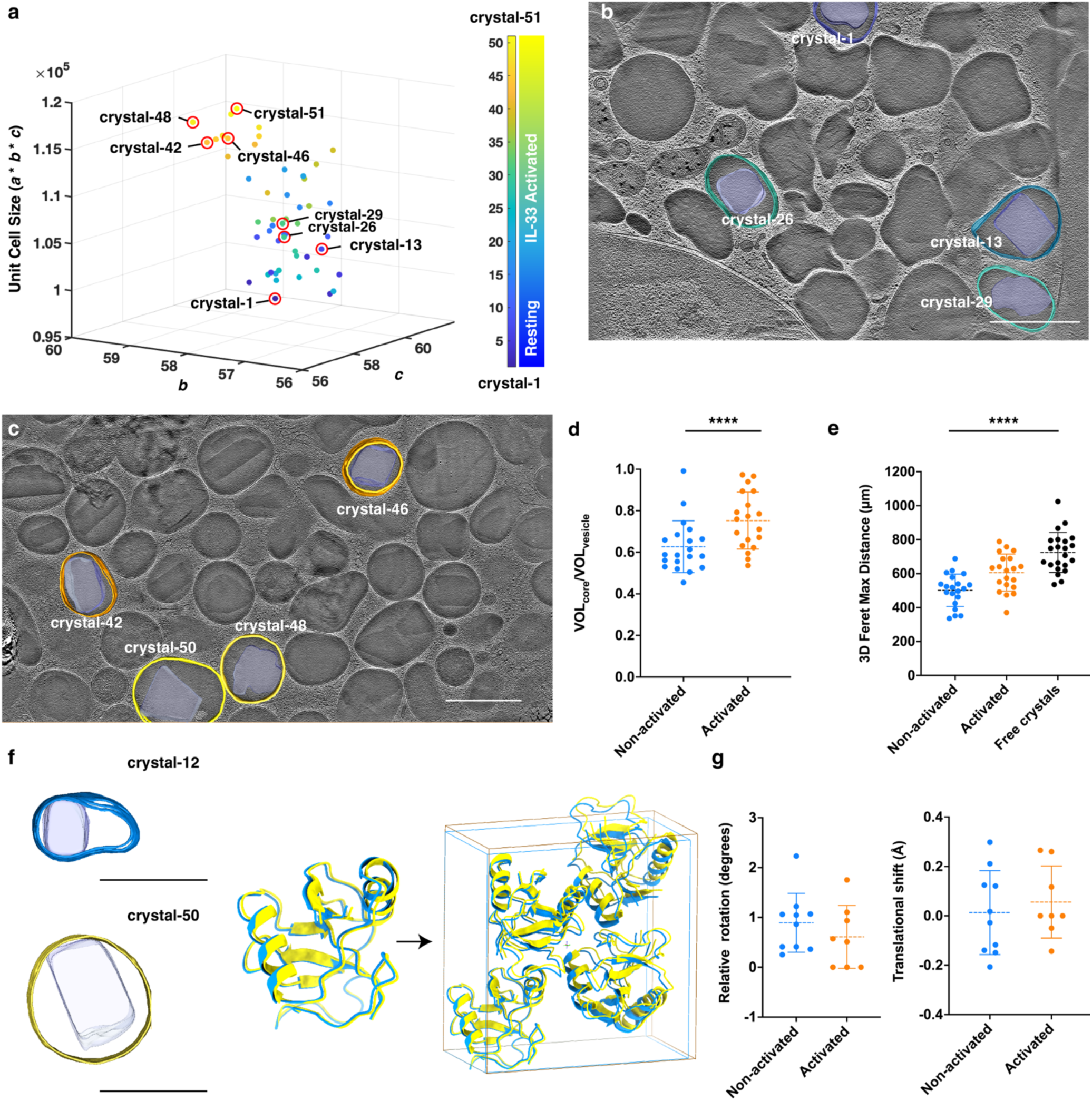
Native-state gMBP-1 nanocrystalline lattice transition upon activation revealed by correlative single-granule profiling. **a**, Unit cell size profiling of individual intragranular crystals (n = 51) from resting (n1 - n20) and IL33-activated human eosinophils (n21 - n51) by MicroED. Unit cell volume (*a* x *b* x *c*, Å^3^) was plotted against the unit cell length on the *b* and *c* axes. Diffracting intracellular granules were indexed starting from resting (n1 = 1) to activated eosinophils (n51 = 51), colored in MATLAB Parula. **b-c**, Diffracting granules were correlated between MicroED and cryo-ET data and were mapped back to their exact locations in the cellular environment. Comparisons of cryo-ET reconstructed volume ratios of crystalline cores to parent granules between non-activated (n = 20) and activated (n = 19) eosinophils (**d**) and 3D maximum Feret distance (**e**) of the crystalline cores from non-activated (n = 20), activated (n = 19) eosinophil cells and membrane-less, free crystals from activated eosinophils (n = 22) by one-way ANOVA test (\*\*\*\**p* < 0.0001). **f**, Changes in unit-cell lattice packing by superimposition of a resting granule (crystal-12) and an IL33-activated granule (crystal-50). **g,** No significant difference existed between the relative rotational and translational shifts of individually refined structures along the 2(*a*) axis within one unit cell. Scale bars = 1 µm in **b, d, f.**

### Intragranular crystal transition and degranulation upon activation

IL33 is a potent activator of eosinophils, triggering degranulation processes^23,37^. We sought to understand the possible mechanisms of how intragranular crystalline cores undergo structural changes upon eosinophil activation. We catalogued and color-coded the raw datasets (n = 51) that were indexed, with crystals 1 through 20 from non-activated eosinophil SGrs, and crystals 21 through 51 from IL33-activated eosinophil SGrs. Plotting the crystal unit cell volumes against the cell axes *b* and *c* indicated a clear trend where the nanocrystals from activated eosinophils had larger unit cell volumes (*a* * *b* * *c*) compared to nanocrystals from non-activated eosinophils (Fig. 2a). Correlated with the real-space information from montage cryo-ET^21^, the larger crystal unit cell volumes (color-coded in the brown to yellow range) were often associated with SGr that carried expanded crystalline cores, and were situated in dynamic cellular environments indicative of active degranulation (Fig. 2c, 3d-g). In contrast, the smaller unit cell size nanocrystals (color-coded in the green to blue to cyan range) were often found in intact SGr where the cores were more compact (Fig. 2b, 3a) relative to granule size.

**Figure 3.**
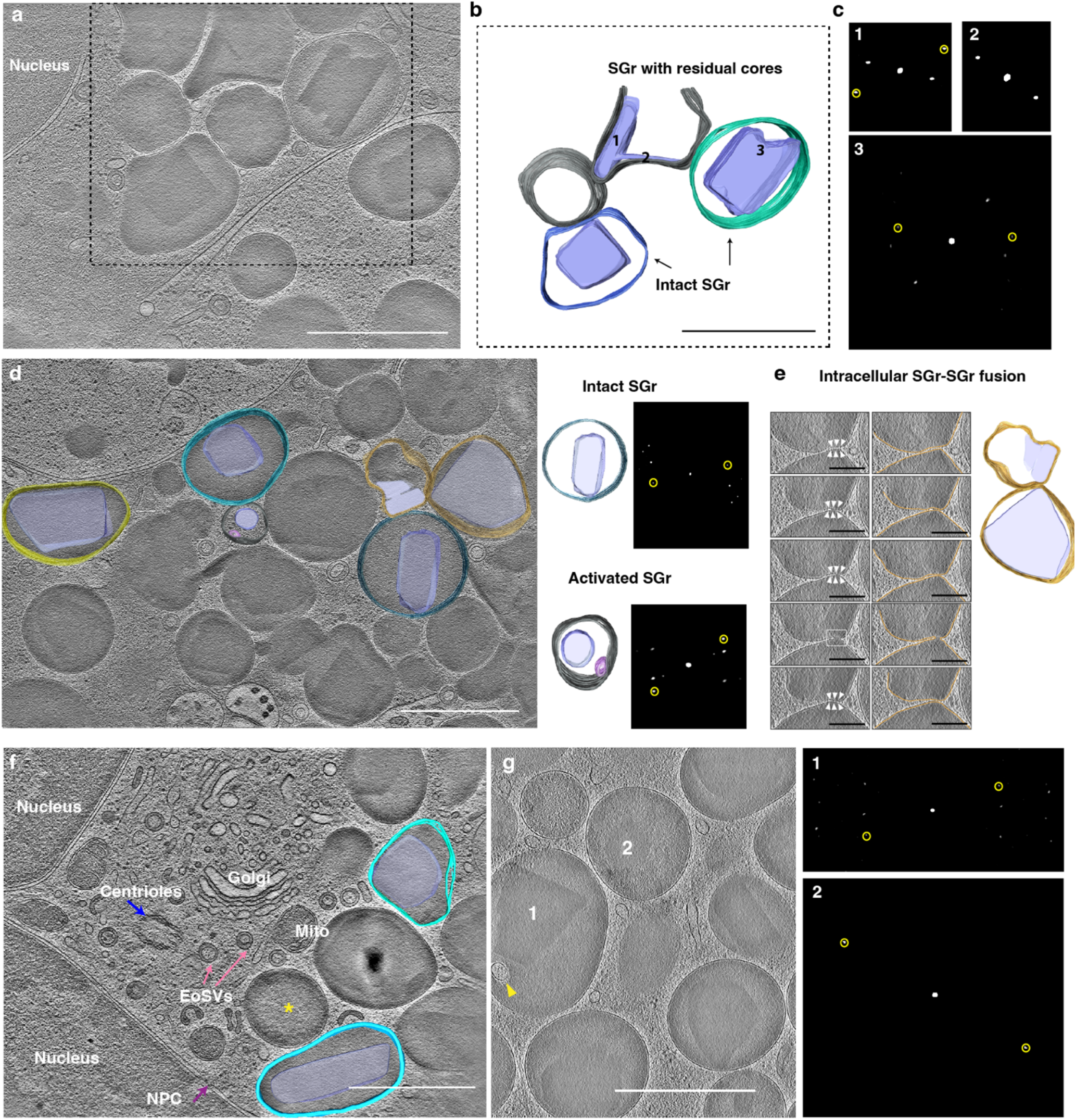
Ultrastructural changes to human eosinophil granules and crystalline cores upon activation. **a-c,** Resting eosinophils contain predominately intact secretory granules (SGr, black arrows, indexed 3) and occasional SGr with residual cores (**a, b,** indexed 1, 2). SGr carry diffracting crystalline structures (indexed 1-3) with corresponding Fast Fourier Transform (FFT) analysis (**c**). **d-g**, IL33-activated eosinophils have SGr of varied morphologies that relate to activation state: intact (**d**), intragranular membrane-containing (**d, g**), empty core (**f**, yellow asterisk), and residual cores (**g**, 2). Diffracting crystals were observed in activated SGr (**d, g**). 3D snapshots of IL33-activated SGr indicated triggering of granule cytokine secretion by piecemeal degranulation (**d, f, g**), and occasional intracellular SGr-SGr fusion (**e**). Sombrero vesicles (EoSVs) (**f**), intragranular membranous structure within activated SGr (**g**, vesicle 1), translucent or empty SGr without cores (yellow asterisk in **f**), residual crystals (**g**, vesicle 2) were frequently seen in IL-33 activated eosinophils. Scale bars of 1 µm in **a, b, d, f, g,** and 200 nm in **e.** FFT analysis were done on the intragranular crystalline with a consistent peak corresponding to 6.28 nm^-1^, circled in yellow.

Consistently, quantitative analyses of crystal volumes relative to SGr vesicle size (Fig. 2d) and crystal dimension (Fig. 2e) showed a significant difference (*p* < 0.005) between the non-activated and IL33 activated granules. Individual granules with reasonable diffraction data completeness (completeness > 50%) were further refined against the final gMBP-1 model (PDB: 9DKZ) using only rigid body and TLS refinement in Phenix^38^ to capture the rearrangement of gMBP-1 monomers within the crystal upon activation. Superimposing individually refined structures and asymmetric unit cells of a representative intragranular crystal-12 from a non-activated eosinophil and crystal-50 from an IL33-activated eosinophil showed a relative rotation of 2° and translational shift of 0.3 Å between two monomers along the 21(*c*)-axis (Fig. 2f). This appeared to contribute to the observed *c*-axis expansion upon activation (Fig. 2a). There was no significant difference in rotation and translation along the two-fold (*a*) axis within one asymmetric unit cell between the non-activated and IL33 activated crystal groups (Fig. 2g), suggesting that the IL33 triggered unit cell growth resulted mainly from crystal packing changes along the two-fold screw axes.

Upon cytokine-mediated activation, eosinophils frequently release their cytotoxic basic proteins via a piecemeal degranulation(PMD) pathway^3^ in which EoSVs, i.e., similar in appearance to the Mexican hat^15^, gather around the SGr and shuttle the protein contents to the extracellular space. SGr-SGr fusion followed by interactions between fused SGrs and the plasma membrane (e.g., compound exocytosis)^39^ is another pathway by which eosinophils secrete granule-derived products. By cryo-ET, the intact SGr in non-activated mature eosinophils displayed a well-defined, condensed, electron-dense nanocrystal core surrounded by the less dense matrix (Fig. 3a-c, d). Consistent with previous reports^15,27^, we saw occasional SGr in non-activated eosinophils that exhibited disassembly of the crystalline core at interfaces with the matrix (Fig. 3b-c, crystal 1 and 2). In IL33-activated eosinophils, SGrs were more heterogeneous. In addition to intact granules (Fig. 3d), internal tubular membranous structures were seen within the granular matrix alongside diffracting crystalline cores, indicative of an early activated state^15,40^ (Fig. 3d, g, crystal-1). The intragranular formation of a membranous network has been suspected of being involved in the formation of EoSVs from SGr^3^. Emptying granules characterized by an electron-lucent core (Fig. 3f, yellow asterisk) and residual crystals (Fig. 3g crystal-2) were noted, and were surrounded by a pool of double-membrane EoSVs with an outer diameter of ∼120 nm, suggesting later stage degranulation^3,15^. Intact, fragmented, or residual crystalloid structures from both non-activated and IL33-activated granules displayed varying levels of lattice packing (Fig. 3c, d, g). Previous studies^15^ suggest that SGr may undergo a gradual process and generate carriers for transportation of granule-derived products including MBP-1^15^. Correlating the expansion of unit cell size, increase in nanocrystal volume relative to the hosting granule (0.62 ± 0.12 versus 0.75 ± 0.13, non-activated versus activated), and various SGr activation states captured within a comprehensive cellular context (Fig. 3d-g) further supported this idea and suggested a potential role for crystal unpacking that allows for the transportation of MBP-1 to the extracellular space. While we saw IL33 predominately trigger PMD processes in activated cells (Fig. 3d, f), montage cryo-ET also captured snapshots of intracellular granule-granule fusion (Fig. 3e) with evidence^39^ of the compound exocytosis degranulation pathway where naked crystal cores were seen budding from cells along the cytoplasmic membrane. This signified a likely fusion of SGr with the plasma membrane to release its granular contents (Extended Data Fig. 5b-f, 6f, g, white arrows).

In summary, IL33 activation triggered the nanocrystal lattice to grow along the two-fold screw axes with the two-fold (*a*)-axis remaining constant (Fig. 3). This implies that the lattice unpacking is directional, mainly driven by molecular interactions along the two-fold screw axes. The nanocrystal disassembly was concurrent with eosinophil degranulation process triggered by cytokine activation.

### Insights into nanocrystal assembly *in-situ*

While eosinophils use various degranulation pathways to transport intragranular mediators to the extracellular space, a conserved observation has been the initiation of intragranular nanocrystal disassembly to solubilize MBP-1. Previous protein crystallography elegantly characterized this very basic protein, MBP-1, under *in vitro* crystallization conditions (pMPB-1, PDB: 1h8u)^18^ and bound with heparin disaccharide (HD) as a complex (PDB: 2brs)^17^. There was a striking difference^18^ between purified then recrystallized MBP-1 (pMBP-1) and other CTL proteins with respect to specific areas of certain domains, particularly the orientation of the long loop L4 (residue 180-191) where calcium-mediates carbohydrate binding in the CTL domains^41–44^. When superimposing *in-situ* SGr MBP-1 (gMBP-1) and purified then recrystallized MBP-1 (pMBP-1), the region with the highest variation, with an RMSD of 0.922 Å across 11 residue pairs compared to the overall RMSD of 0.623 Å across 111 residue pairs, also resides in this calcium-mediated carbohydrate binding pocket (Fig. 4a-b). In the structure of gMBP-1, Trp185-Pro190-Trp191 adopt a downward pointing pocket configuration that stabilizes Pro190 with a series of aromatic residues (Trp191, Trp185, Tyr184, Phe182, Trp180), compared to an arched, outward pointing loop observed with recrystallized MBP-1 (Fig. 4a-b). In the recrystallized MBP-1 (pMBP-1), this arched configuration shifts residues Arg193, His196, Arg208, Arg209 forward to interface with sulfated heparin disaccharide ^17,18^, thus neutralizing the charge of the binding pocket (Fig. 4a-b). In contrast, the downward pointing pocket configuration adopted by the gMBP-1 binding region allows for the placement of Pro190 into a neutralized and hydrophobic cavity that promotes favorable interactions^45^ between proline and surrounding aromatic residues^46^ (Fig. 4c-f). It has been found that aromatic-proline sequences more readily assume cis-prolyl amide bonds^46^. Consistently, in gMBP-1, the aromatic rings (Trp185 and Trp191) interact with the ring of Pro190 in its cis conformation. As a result, the nearby hydrophilic groove created by Ala186-Ala187-His188-Gln189 (Fig. 4c-f, blue ribbon) accommodates Arg130 of the neighboring of gMBP-1 monomer (Fig. 4d-f, grey ribbon) positioned along the 21(*c*)-axis. Furthermore, the aromatic residues Tyr129 and Try222 of the neighboring monomer (Fig. 4d, grey ribbon) contribute to the enrichment of aromatic side chains in the pocket region, stabilizing the local aromatic-proline interactions. Thus, purified recrystallized MBP1 (pMBP-1) rather than granule MBP-1 (gMBP-1) is the outlier in not sharing the conserved, downward pointing pocket loop configuration of the other CTLs with the proline (Pro190 in gMPB-1) in its cis-form^41–44^ (Extended Data Fig. 11a-c).

**Figure 4.**
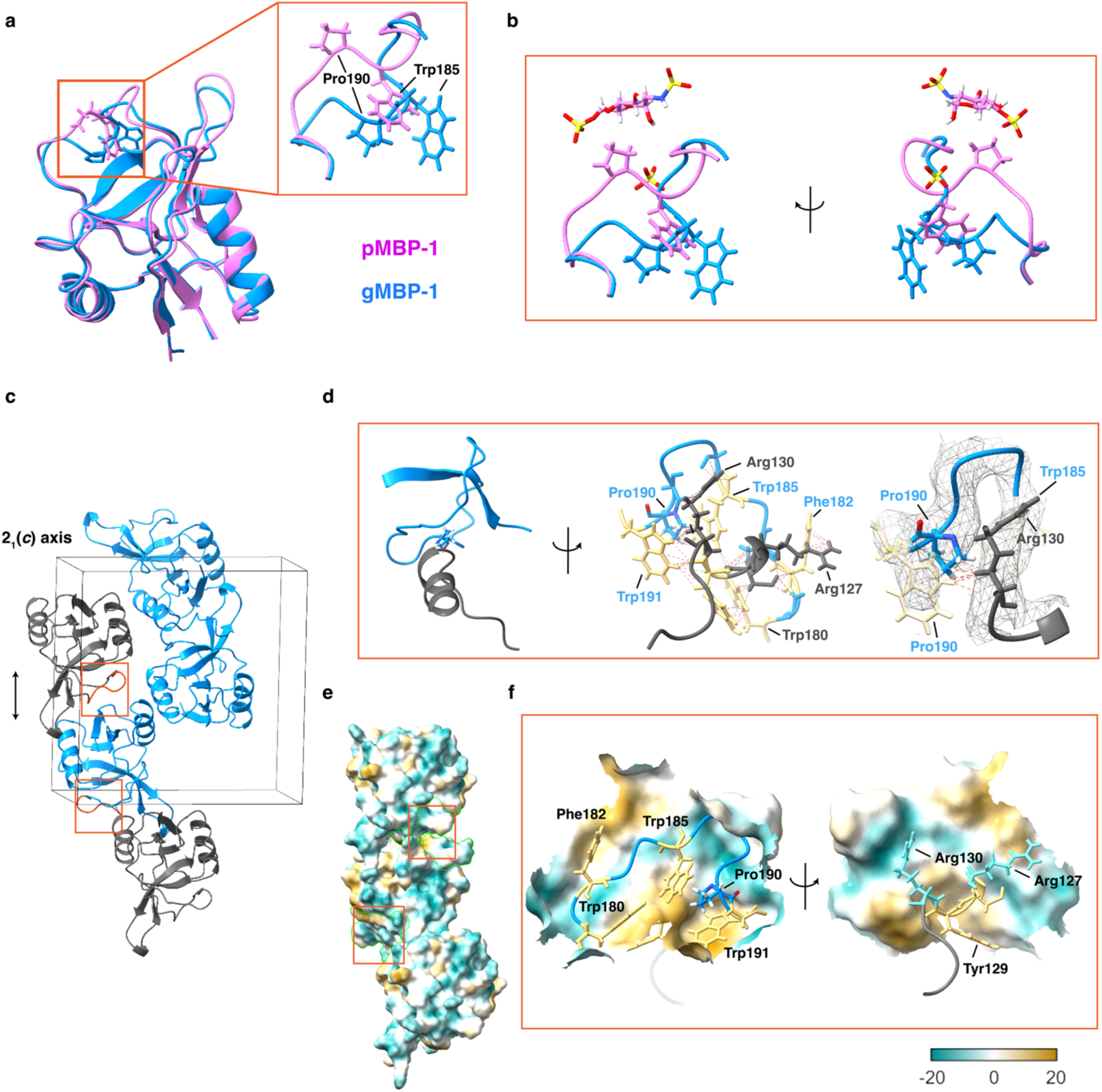
Distinct structural difference in the canonical calcium binding pocket regions of *in-situ* and *in-vitro* crystal forms of human eosinophil major basic protein-1 (MBP-1). **a-b,** The loop region (Residue 180 to 191) that accommodates heparin disaccharide (HD)/sulfate binding in *in-vitro* MBP-1 displayed structural differences in both residue configuration and the carbon backbone when *in-situ* granular MBP-1 (gMBP-1, PDB: 9DKZ, blue) was superimposed on *in-vitro* purified and recrystallized MBP-1 (pMPB-1, PDB: 2BRS, pink). **c,** The *in-situ* crystal packing of gMBP-1 in one unit cell with the carbohydrate binding loop region highlighted and boxed in orange. The monomers along the 21 (*c*) axis are alternatingly colored in blue and grey. **d-f,** The detailed residues (**d**) and calculated molecular lipophilicity potential surface (**e, f,** dark cyan for the most hydrophilic, white and dark goldenrod for the most hydrophobic, on a scale range of -20 to 20) views of the binding-pocket loop region between two monomers (blue and grey) along the 2_1_(*c*) axis. The loop configuration supports the preferrable interaction of Pro190 with the surrounding aromatic residues (Tyr129, 184, 222, and Trp180, 185, 191, and Phe182 are highlighted in yellow) from the two monomers: Trp180, 185, 191, Phe182, and Try 184 from blue monomer, Try129 and 222 from the neighboring grey monomer on the 2_1_ (*c*) axis). The residues are labeled corresponding to the monomer color.

This conserved loop configuration places the hydrogens on the pyrrolidine ring of the proline (Pro190 in gMBP-1) perpendicular to the indole ring of tryptophan (Trp185 in gMBP-1) five residue away (Extended Data Fig. 11b). Both residues participate in defining the CTL fold. Sequence comparisons of the CTL domain across eosinophil MBP-1 homologues indicate the conservation of the tryptophan and proline in this loop region, except for the chimpanzee that has a leucine (Leu190) in this corresponding position (pPRG2, Extended Data Fig. 11d). Despite this difference, Alphafold model of pPRG2^47^ predicts a similar configuration of Leu190, forming a pocket and overall downward pointing loop architecture (Extended Data Fig. 11c).

To further define nanocrystal formation *in-situ*, we compared our gMBP-1 with the recently resolved cryo-EM structure of proMBP-1^48^ bound to the metalloproteinase pregnancy-associated protein A (PAPP-A). ProMBP-1 is the unprocessed form of mature MBP-1 and consists of an acidic N-terminal propiece followed by a highly basic CTL domain, namely MBP-1, at the C-terminus (Extended Data Fig. 12a). In the cryo-EM structure of proMBP-1^48^, the propiece (residue 17 to 106) is resolved from residue 88 to 106. Close inspection shows that proMBP-1 adopts the downward pointing loop configuration, allowing for proper positioning of aromatic residues Trp180, Phe182, Tyr184, Trp185, and Trp191 to interface with Pro190 and multiple prolines (Pro692, 693, 696) of PAPP-A in the hydrophobic groove (Extended Data Fig. 12b). We superimposed two proMBP-1 molecules along the 21(*c*)-axis on two gMBP-1 monomers within one unit cell. Glutamate-rich residue 88 to 92 are predicted to interfere sterically with the monomer-monomer interactions on the *c*-axis, thus hindering the formation of crystal lattice.

These results demonstrate that the downward pointing loop configuration in the canonical calcium binding pocket and intra- and inter MBP-1 monomer-monomer interactions along the 21(*c*) axis play a crucial role in intragranular nanocrystal formation, stabilization, and mobilization.

## Discussion

Quantitative electron microscopy^3^ and functional analyses^49^ of human eosinophils indicate that human eosinophil major basic protein-1 (MBP-1) forms intragranular nanocrystals that are transported to extracellular space upon cell activation^11^, thereby participating in immune-response activities. The well-organized crystal cores in mature secretory granules (SGr) are thought to function as inert storage bodies to protect eosinophils from the non-selective cytotoxicity of MBP-1^10^. This hypothesis is supported by discovery of the pre-pro form of MBP-1 (proMBP-1)^8^ and functional characterization^9,10^ of MPB-1 self-assembly and dissolving under environments associated with immunopathogenic diseases. A major accomplishment of our work is the determination of the *in-situ* structure of granule MBP-1(gMBP-1). Using cryo-FIB milling, MicroED, and cryo-ET, we probed nanocrystal cores directly in unperturbed SGr situated inside donor-derived eosinophils. The native crystal lattice packing and structure of gMBP-1 presented here provide further details about *in-situ* intragranular nanocrystal formation and new insight into the degranulation process in the context of the cellular landscape from montage cryo-ET.

One of our most notable findings is the configuration of the loop region of granule MBP-1 (gMBP-1) that corresponds with the calcium-mediated carbohydrate binding site of other canonical CTL domain proteins. In the CTL family^50^, one of the calcium binding sites is in the L3-L4 region comprised of highly conserved acidic residues such as Glu80, Asn82, Asn83, Asn105, Asn106, Glu107 in E-Selectin^42^ (Extended Fig. 11c). First, while gMBP-1 has a similar overall topology of CTL domains, as pointed out by Swaminathan et al^18^., it does not share those conserved functional residues. The equivalent MBP-1 residues to this binding site region are Gln189, Trp191, Ser192, Arg208, Arg209, Ala210. None of them are acidic, thus effectively abolishing the ability of gMBP-1 to coordinate calcium. Second, in purified and recrystallized MBP-1 (pMBP-1), this calcium-carbohydrate binding site has a distinct structure (Fig. 4a, ribbon representation in pink), with the L4 loop region pointing up versus pointing down when compared to other CTL proteins^18^. Through the pointing up configuration, *in vitro* pMBP-1 interacts with sulfated sugars such as heparin or heparin sulfate^17^ (Fig. 4a), suggesting that the site adapts to bind different carbohydrate ligands by a mechanism that does not require the coordination of calcium. Our results prove that in contrast to recrystallized MBP-1, granule MBP-1 adopts a similar downward pointing loop architecture in this critical pocket, just like those present in the canonical CTL domains^41–43^ (Fig. 4a, Extended Data Fig. 11a-c).

Importantly, rather than coordinating with calcium, this downward pointing configuration of gMBP-1 is architecturally stabilized by favorable cis-proline-aromatic interactions (Pro190 in gMBP-1) with residues from two monomers positioned along the 21(*c*) axis within one unit cell. We propose that favored intra- and inter-molecular interactions between the cis-proline in the downward pointing loop pocket and aromatic residues are crucial for initializing and stabilizing the nanocrystal within SGr.

Our hypothesis is further supported by investigations of proMBP-1, the nontoxic precursor of mature MPB-1^8^. proMBP-1 consists of an acidic propiece (residues 17-106) and basic MBP-1 protein (residues 107-222). The propiece is believed to neutralize the basic nature of MBP-1 and protect the cell during transport from Golgi apparatus to the granule in eosinophils^8,10^. The propiece residues pack against one side of MBP-1 to extend the β1-β7 sheet before making a sharp bend that harbors cleavage sites between the propiece and MBP-1. This bend is stabilized by a disulfide bond (Cys104-107) that must be broken to release the propiece from MBP-1^51^. proMPB-1/PRG2 is soluble and functions to inhibit proteolytic activity of pregnancy-associated plasma protein A (PAPP-A) by complex formation^51^. A recent cryo-EM structure^48^ of proMBP-1 in complex with PAPP-A revealed that one of the proMBP-1-PAPP-A interfaces is located within the same canonical calcium-mediated carbohydrate binding region. Unsurprisingly, proMBP-1 adopts downward pointing loop configuration, allowing for proper positioning of aromatic residues Trp180, Phe182, Tyr184, Trp185, and Trp191 to interface with Pro190 of MBP-1 domain and multiple prolines (Pro692, 693, 696) of PAPP-A in the hydrophobic groove formed in this macromolecular complex (Extended Data Fig. 12a).

Docking proMBP-1 monomers in the crystal lattice of gMBP-1 demonstrate that cleavage of its pro-domain is a prerequisite for intragranular nanocrystal formation in mature SGr of human eosinophils. In addition, the must-be-cleaved disulfide Cys104-107 in the propiece would sterically interfere with the arginine-rich (Arg168-Arg170) region of the flexible long loop of the neighboring proMBP-1 monomer along the 21(*c*) axis. This might explain why this flexible long loop region was poorly resolved in our *in-situ* granule MBP-1 (gMBP-1) structure. Together, these analyses support our proposal that, in the case of gMBP-1, the canonical calcium-binding pocket of CTL domain and associated downward pointing loop configuration directly contribute to intragranular nanocrystal formation by driving intermolecular interactions on the two-fold screw axis along *c*, which strengthens the concept that successful crystallization of MBP-1 is critical for its nontoxic storage in human eosinophil SGr. This is consistent with the observation that IL33-activation triggered the nanocrystal unit cell expansion along the 21(*c*)- axis, during degranulation (Fig. 3a, f).

How do intragranular nanocrystals dissemble during degranulation? Previous studies by numerous investigators^3^ have provided a rich spectrum of activated SGr states, including granule enlargement, core disarrangement, and content loss. These are suggestive of a dynamic process that involves changes in crystal lattice unpacking. We took advantage of cryogenic sample preparation that preserves molecules in a near native state^52^, and further characterized individual granules in their unperturbed cellular context via montage parallel array cryo-tomography (MPACT)^21^. Correlating the real space 3D volume information obtained from MPACT and diffraction space lattice arrangement from MicroED at a single granule level (Fig. 3-4, Extended Data Fig. 7), we observed a direct link between a directional growth along the two-fold screw axes of the unit cell and volume expansion of the intragranular core relative to the hosting SGr (Fig. 2d-e). Reciprocal space analysis via Fast Fourier Transform (FFT) of nanocrystal tomograms showed consistent peaks corresponding to 6.28 nm^-1^ in both non-activated and activated SGrs (Fig. 3c, d, g). We do not know which lattice space dimension corresponds to 6.28 nm^-1^ due to the random orientations of the intragranular crystals and similar unit cell lengths of *b* and *c* (57.87 Å and 59.06 Å, respectively), which are twice the cell length of *a* (*a* = 31.24 Å). Despite that, the concurrence of SGr intragranular membranous structures adjacent to the nanocrystal core, and a pool of SGr nearby or attached to EoSVs, implies that a process of MBP-1 active transport may be coordinated by MBP-1 crystal disassembly from the SGr to the extracellular space. Early studies^16^ show that eosinophil SGr have lysosomal propensities that enable acidification to occur in the intragranular matrix upon activation. The current resolution of gMBP-1 (3.2 Å) did not provide enough information for investigation of per-residue protonation, especially around the downward pointing loop pocket. However, the correlative results from both real-space and diffraction analyses allow us to propose that nanocrystal packing of gMBP-1 is mediated by the intra-and inter cis-proline-aromatic monomer-monomer interactions that becomes unstable, likely due to intragranular acidification upon activation, thus leading to crystal disassembly along the two-fold screw axes and release of MBP-1 from its nanocrystal form. The “soluble” MBP-1 proteins are ready for uptake by intragranular membranous structures, followed by transport to extracellular space via EoSVs.

Unexpectedly, we saw membrane-less, free crystals in proximity of or seemingly in the cytoplasm of IL33-activated eosinophils, especially in flattened cell membrane extensions that were distinctly separated from the main cell body where SGrs were densely packed. These were not observed in cryo-EM of non-activated eosinophils (Extended Data Fig. 2). Previous work^10^ showed that the toxicity of MBP-1 is closely associated with its aggregation state. While MBP-1 aggregates to form prefibrillar and fibrillar oligomers that cause damage to mammalian cells, mature fibrils or heparin-enhanced aggregates are toxicity neutralized^10^. While free crystals were unable to diffract sufficiently for structure determination, clear crystal lattices were observed (Extended Data Fig. 3d-f), suggesting a difference in MBP-1 maturity and a possible lower cellular toxicity level. It is possible that IL33-activation triggers the expansion of the cell edge and reorganization of the SGr to create flattened extended areas. As a result, clustered SGr could release less toxic nanocrystals into these sheltered regions, similar to eosinophil extracellular traps (EETs)^53^. EETs are enriched in released MBP-1 and mitochondria and nucleus DNA within a relatively concentrated zone to immobilize and kill pathogens.

Curiously, membrane-less free crystals from activated eosinophils had the largest size dimension when compared to intragranular nanocrystals (Fig. 3e). SGrs share lysosomal characteristics and becomes acidic upon activation. Crystals and protein aggregates^31^ are also known to increase the permeability of lysosomal membranes that eventually cause them to rupture. The delimiting SGr membrane may become ruptured towards late activation states when the intragranular nanocrystal expands in size. Future studies of this enclosed microenvironment are required to understand how these free crystals are formed, and whether the extrusion of free crystals is unique to IL33-activated eosinophils.

How does purified and recrystallized MBP-1, specifically residues 185-191, adopt the opposite upward pointing configuration in the critical L4 loop region? Heparin, a known enhancer of MBP-1 aggregation, irreversibly binds free MBP-1 and promotes its recrystallization^17,18^. We speculate that loop flipping could be a result of crystal disassembly. Alternatively, heparin binding could trigger the configuration change of soluble MBP-1, followed by its recrystallization. Site directed mutagenesis within the loop region could help elucidate the flipping mechanism. However, the *in vitro* experiments might alter other interfaces that stabilize overall protein structure or diminish functional relevance of MBP-1. Other experimental options could be explored to assess gMBP-1 activation and function over a varied pH range using molecular dynamics simulations^54^.

Here, we developed and applied an integrated *in-situ* cryo-EM workflow to interrogate the structure of protein nanocrystals within human cells, specifically eosinophils. This investigation marks one of the first *in-situ* structural reports on human protein nanocrystals. Through this study, we determined the *in-situ* structure of human intragranular eosinophil major basic protein-1 (gMBP-1) to define mechanistic processes associated with eosinophil degranulation upon cytokine activation. MicroED^20^ enabled atomic structure determination of micrometer-sized 3D nanocrystals. Cryo-FIB milling^19^ and cryo-ET, including montage cryo-ET^21^, supported the fabrication of lamellae and the analysis of macromolecules in the large expanse of cell interiors. Along with increasing interests in *in vivo* protein crystallization^55^, this correlative microscopy framework will be broadly applicable to native environment structural studies of human insulin crystals^56^, virus-induced cholesterol crystals^57^, and other multi-phasic macromolecules implicated in human health.

## Methods

### Donor-derived peripheral blood eosinophil isolation, activation, and viability measurements

Eosinophils were from donors with atopy (**Supplemental Table 1**) who had eosinophil counts in the normal range of 200−500 per μL. The studies were approved by the University of Wisconsin-Madison Health Sciences Institutional Review Board (protocol No. 2013–1570). Informed written consent was obtained before participation. On the day of experimentation, eosinophils were purified from 200 mL of heparinized blood from a single donor. Centrifugation through a Percoll (density of 1.090 g/mL, Sigma-Aldrich) gradient separated eosinophilic and neutrophilic granulocytes from mononuclear cells. Negative selection using the AutoMACS system (Miltenyi, Auburn, CA) and a cocktail of anti-CD16-, anti-CD14-, anti-CD3-, and antiglycophorin A-coupled magnetic beads removed neutrophils and red blood cell precursors. Purity of eosinophils compared to other leukocytes was ≥98% as determined by Wright−Giemsa staining followed by microscopic scoring of cells. To allow recovery from purification, eosinophils (10^6^ per mL) were placed in 1640 RPMI medium (Sigma-Aldrich) supplemented with 0.1% human serum albumin (HSA) (Sigma-Aldrich) for 2 hr at 37℃ in a 5% CO2 incubator prior to further experimental treatments. Viability of eosinophils was determined using a Live/Dead cell viability kit (80 nM Calcein-AM, 400 nM ethidium homodimer-1, Cat. L3224, ThermoFisher Scientific, USA) either of cells in suspension or after addition to TEM grids. For fluorescently stained suspended eosinophils, glass-bottomed culture dishes (35 mm dish /20 mm glass diameter, Cat. P35G-1.5-20-C, MatTek Corp., MA, USA) were used to disperse the cells for 5 min prior to live-cell fluorescence imaging. Isolated non-activated eosinophil viability was > 99%. Occasional cell death was observed in IL33-activated eosinophils when the activation time was more than 1 hour.

### Electron microscopy grid preparation, live-cell fluorescence microscopy, and vitrification

To prepare eosinophils (non-activated or IL33-activated) samples for cryo-EM, 200 mesh gold Quantifoil R1.2/20 grids (Quantifoil, Germany) were glow discharged for 60 sec at 10 mA and then soaked in 70% (v/v) ethanol for >20 min. The grids were washed three times in cell-culture grade water (Cat. 25-055-CVC, Corning. Corp. NY, USA) and PBS pH 7.4 (Cat. 10010023, ThermoFisher Scientific, USA), followed by an application of fibrinogen (10 µg/mL) and incubation in a 5% CO2 tissue culture incubator at 37℃ for >2 hours. Following fibrinogen coating, the grids were washed three times with RPMI-1640 supplemented with 0.1% HSA After the final wash, eosinophils were applied to the grids. The TEM grids with non-activated eosinophils were directly plunge frozen after approximately 5-10 min of incubation. The second set of rested cells applied to the TEM grids were allowed to settle on the EM grids for approximately 5-10 min and were then activated with IL33 (50 ng/mL, R and D Systems, Minneapolis, MN) for 1 hr at 37℃, prior to plunge freezing. For fluorescent staining of membranes, non-activated or IL33 activated (after 40 min of activation) eosinophils were incubated with low-toxicity CellBrite Steady Red cytoplasmic membrane dye (1:200 dilution, Ex/Em 562/579 nm, Cat. 30107-T, Biotium, CA, USA) at 37℃ for 20 min. Alternatively, we used CellMask (1:1000 dilution, Ex/Em 649/666 nm, Cat. C10046, Invitrogen, USA) to label the plasma membrane (37℃ for 5 min). CellBrite performed better as to longer dye retention on the plasma membrane versus internalization, and low toxicity to eosinophil. However, to ensure native preservation and minimal disturbance of intragranular crystals, the structural determination by MicroED and MPACT was conducted on unstained or non-labeled eosinophils. The fluorescently stained cells on the grids were then gently washed two times with RPMI-1640 supplemented with 0.1% HSA. The TEM grids with the cells were placed on the glass bottom of a MatTek dish (MatTek Corp., MA, USA ), were examined via live-cell imaging at 20x magnification (0.4 NA lens, dry) and 63x magnification (1.4 NA lens, oil-immersion) in brightfield and filter cubes of FITC (emission, *λ* = 527/30 nm) and TXR (emission *γ* = 630/75 nm) using a Leica DMi8 system (Leica Microsystems, Germany). The grids with non-activated or IL33-activated (in total of 1 hr activation) human eosinophils were vitrified in liquid ethane using a Leica EM GP 1 (Leica Microsystems, Germany). The Leica EM GP 1 plunger was set to 37 ℃, 95% humidity, and blot time of 8-10 sec for single-sided back-side blotting and plunge freezing. Plunge-frozen grids were then clipped into Autogrids (ThermoFisher Scientific) and stored in cryo-grid boxes under liquid nitrogen.

### Single-granule profiling

The single-granule profiling workflow consisted of five main steps: (1) cell identification via low-dose 2D cryo-EM, (2) cryo-FIB milling, (3) granule identification, (4) per-granule diffraction acquisition via MicroED, (5) per-granule montage cryo-ET via MPACT. Detailed procedures and experimental conditions for each step are described in the following individual sections. Frist, non-activated or IL33-activated human eosinophils were identified by 2D cryo-EM imaging under low dose conditions based on morphologic characterization following the established conventions. Eosinophils can be uniquely distinguished by their morphological shape. Non-activated human eosinophils are spherical with irregular short surface protrusions. Stimulated or activated eosinophils appear as more flattened and elongated cells^3^. The clipped grids were loaded onto a Titan Krios G4 (ThermoFisher Scientific) operated at 300 kV in EFTEM mode. Using SerialEM (Version 4.0.10), grid overview maps (EFTEM, 125 x, 1007 Å/pixel) and square view maps (EFTEM, 125x, 399 Å/pixel) were collected to identify non-activated or IL33-activated eosinophils on TEM grids, respectively. The total dose was neglectable at ∼0 e/Å^2^. Second, the pre-screened grids with targeted non-activated or IL33-activated eosinophils were loaded onto a cryo-FIB-SEM dual beam system Aquilos 2 (ThermoFisher Scientific). Cells identified by low dose cryo-EM imaging in step 1 were cryo-FIB milled close to the center of the cell bodies to generate the final 200-250 nm lamellae.

Third, after loading the grids containing multiple lamellae on the Titan Krios G4, low dose grid overview images (EFTEM, 125 x, 1007 Å/pixel) and SA-magnification (EFTEM, 38.52 Å/pixel, defocus of -100 µm, slit width of 10 eV) image montages were acquired to capture whole individual lamella at the correct pre-tilt angles. The dose per image frame in the EFTEM SA magnification imaging mode was <0.01 e/Å^2^. Most of the areas were only imaged once at the SA mapping condition with a 10% overlap between image montage tiles. The low dose SA-montages of the lamellae were used to mark possible secretory granules (e.g., dark vesicles with condensed central densities) for subsequent MicroED acquisition. Fourth, diffraction data sets were collected of each marked granule following a continuous stage rotation MicroED acquisition scheme using TEM mode, with the beam stop inserted, on a Ceta-D camera (ThermoFisher Scientific). Fifth, after all MicroED data sets were collected of the marked granules, each granule was revisited for montage parallel array cryo-tomography (MPACT) collection using EFTEM mode on the Falcon 4i with SelectrisX energy filter (ThermoFisher Scientific). As a result, montage tomograms and MicroED data sets were correlated to create per-cell and per-granule profiles. Diffraction data were used to determine the *in-situ* structure and unit-cell dimensions of mature human eosinophil major basic protein-1 (gMBP-1). Tomography data were used to examine the shape, relative volume, and unperturbed granular environment of granule crystals in three dimensions (3D). This workflow allowed for correlation of real space and diffraction data at a single cell and single granule level.

### Cryo-focused ion-beam milling

Cryo-FIB milling of the eosinophils was performed following the previously published protocols^58^. After initial screening by cryo-TEM to identify non-activated or IL33-activated eosinophils, the clipped grids were loaded into a cryo-FIB-SEM dual beam Aquilos 2 system (ThermoFisher Scientific) operated under cryogenic conditions (stage and shield temperature of ∼192 ℃). Cryo-SEM (2 kV, 25 pA) grid overview images were taken both prior to and after platinum sputtering (20 mA, 25 sec, thickness of 30-50 nm) and organometallic platinum (in-chamber gas injection system, GIS) coating (thickness of ∼3 µm). Milling sites were set up for automated fabrication in MAPS (Version 3.25, ThermoFisher Scientific) and AutoTEM (Version 2.4, ThermoFisher Scientific) with a shallow FIB-milling angle of 8-12° on a 35° pre-tilt Autogrid shuttle (ThermoFisher Scientific). Micro-expansion joints/trenches (width of 500 nm)^59^ were generated using 0.3 nA (rectangular milling pattern, dwell time of 1 µs) around the lamellae sites (usually 10-14 µm in width) to minimize lamellae bending or cracking. The ion-beam milling process was performed using 0.3 nA for rough milling with gradually decreasing currents of 0.1 nA, 50 pA, 30 pA, and 10 pA. The milling time was adjusted based on the size of each cell. The final targeted thickness for the lamellae was 200-250 nm. Prior to unloading, a very thin layer of platinum (20 mA, 3 sec) was sputtered onto the surface of the sample. The preclipped grids with FIB-fabricated lamellae were then loaded onto a Titan Krios G4 (ThermoFisher Scientific) for MicroED and MPACT collection.

### Micro-electron diffraction data collection

The lamellae were rotated 90° relative to the milling direction and loaded onto a Titan Krios G4 operated at -190 ℃ with an accelerating voltage of 300 kV (∼0.0197 Å wavelength). SerialEM^60^ (version 4.0.10) was used to collect 2D montage overview images of each lamella at the correct pre-tilt angles using an EFTEM magnification of 6500x (38.52 Å/pixel, defocus of -100 µm, slit width of 10 eV) in a low dose imaging state of <0.01 e/Å^2^ per frame. Secretory granule sites (x and y stage coordinates) were identified on the 2D lamellae montage views and added to a SerialEM Navigator file. Granule sites were grouped based on their proximity to one another, and eucentricity (z stage coordinate) was refined and updated per group via SerialEM. A new batch session was set up in the EPUD software (Version 1.13. ThermoFisher Scientific) and the XYZ coordinate information per granule from SerialEM Navigator (StagePosition) was then imported into EPUD as batch collection sites by modification of the session xml file. The microscope was switched to TEM mode for MicroED acquisition using EPUD and a CetaD detector. The sample-to-detector distance was set to a calibrated distance of 3454 mm. The diffraction data were acquired by continuously rotating the stage at a rate of 1°/s with a 1°/ tilt angle increment to cover a total tilt range of 40° around the pre-tilt angle calculated from the milling angle (8-12°) and sample loading orientation on the TEM. Low current density conditions were used with a 50 µm C2 aperture and spot size of 11 (gun lens of 1) to generate an illuminated area of 1.28 µm. Based on previous reports^34^, the current density under this imaging condition was 0.15 or 0.2 e/ Å^2^/s. The critical dose was determined by collecting a static diffraction frame each granule at the pre-tilt angle prior to and after a full tilt series of 40°. The normalized intensity level of four diffracted spots ranging from resolutions of 3 to 7° was compared to determine dose effect (n = 3) using the two static diffraction frames. Using the same cutoff reported previously^34^, the critical total tolerable dose was determined when the average of the normalized intensity of the spots on the post-tilt series diffracted frame dropped below 1. A beam stop was inserted during diffraction data set acquisitions. A total accumulated dose of 6.5-8 e/Å^2^ applied to each granule crystal.

### Correlative montage parallel array cryo-tomography (MPACT) data collection and processing

Montage cryo-ET via MPACT was acquired, following previously published conditions^21^ and summarized in Supplemental Table 2. After acquiring diffraction data sets of the secretory granules, the TEM microscope was returned to EFTEM mode (10 eV slit) and operated at 300 kV without changing the gun lens (gun lens of 1). Based on the 2D image montages of the lamellae where diffraction data sets had been acquired, 2×2 or 3×3 MPACT tilt series were set up with the pre-tilt conditions needed for each lamella using SerialEM (Version 4.0.10). The acquisition magnification was EFTEM 26000x (pixel size of 4.727 Å/pixel). A dose symmetric scheme with 3° increments, groups of 2 tilts, and a nominal defocus of -5 µm was applied. The 4096 x 4096 pixels raw frame images were collected and saved in the Electron Event Representation (EER) format. Additional parameters for MPACT acquisition included 12% overlaps of tile frames in both X and Y directions and default spiral setting (Afinal = 1.5, Period = 3, Turns = 50, Revolution = 15). A series of static images, e.g., dose series, over several (n = 2) post-MicroED marked granules were collected until obvious beam damage to the crystalline cores were observed. The tolerated accumulated dose was 100-110 e/Å^2^. The total dose per MPACT tile tilt series was calculated to be ∼50% of the dose series test (e.g., 50-55 e/Å^2^). After MPACT acquisition, all raw EER movie frames were grouped into fractions and brought to 8K super-resolution for alignment and motion correction via MotionCor2^61^. Individual motion-corrected tile frames were stitched per tilt, followed by assembly of a stitched full tilt series using an automated pre-processing pipeline^21^. The stitched tilt series were then binned by 2 (pixel size of 9.454 Å/pixel), aligned via patch tracking, and reconstructed via weighted back projection using IMOD^62^ (Version 4.12). Raw reconstructed tomograms were further processed using IsoNet^63^ at pixel size of 18.908 Å/pixel to reduce the missing wedge effect and improve the signal-to-noise ratio. Briefly, five MPACT reconstructed tomograms were used for IsoNet training with a total of 55 iterations. The trained model was then applied to all MPACT tomograms. Raw unfiltered tomograms (pixel size of 9.454 Å/pixel) were used to identify lattice packing via Fast Fourier Transform (FFT) analysis, while post-IsoNet tomograms were used for visualization, segmentation, and data interpretation. Segmentation was performed in DragonFly (Version 2022.2 Build 1399, Object Research Systems, Comet Group) using the Deep Learning module following procedures reported previously^64^. The automated segmentation results were exported as Tiff stacks and imported into Amira 3D (Version 2023.2, ThermoFisher Scientific). The segmentations of secretory granule limiting membranes and cores were manually cleaned, and Surface Area Volume analysis was performed in Amira 3D. The relative volume (VOLcore/VOLvesicle) of each secretory granule was calculated and compared between non-activated and IL33 activated eosinophil cells using Student’s T-test. Prism (Version 10.1.0, GraphPad) was used to plot the graphs.

### MicroED data processing, refinement, model building, and validation

The diffraction frames were recorded in the standard X-ray diffraction SMV image file format (.img) by EPUD. The diffraction data sets from both non-activated and IL33-activated eosinophil granules were indexed and integrated in XDS^65^. Empty frames were excluded from integration. The maximum resolution cutoff per data set was based on I/Sigma (>0.8). In total, 51 diffraction data sets were reliably indexed. BLEND^66^ was used to guide merging based on cluster analysis with unmerged HKL files of individual data sets as inputs. Dendrogram and absolute equivalence values (aLCV) were used to determine the isomorphism between data sets. The data sets were then scaled using XSCALE and merged based on the BLEND^66^ cluster (aLCV < 1.5) for further examination using post-correction cross-correlation between input data sets (Initial cutoff of correlation between i, j > 0.7). Molecular replacement was carried out in Phaser^36^ from Phenix (v.1.21.1-5286), using an *in vitro* crystal structure of the MBP-1 monomer (PDB: 1h8u.pbd1)^18^. Alternatively, an AlphaFold^47^ model predicted by the AlphaFold Model Prediction module in Phenix^67^ was also used for initial phase determination using the input sequence (1h8u.fa) via molecular replacement and Phaser. Iterative rounds of structural refinement and model building were carried out in phenix.refine^38^ and Coot^68^. To minimize model bias, the composite omit map^69^ was generated using phenix.composit_omit_map in the Phenix program suite, showing a good agreement with the original map. It indicated that the MR phase solution was high-quality and not dominated by model bias. The final structure (PDB code: 9DKZ) of the *in-situ* form of the human major basic protein-1 was merged from 6 diffraction data sets of rested eosinophil granule cores with good cross-correlation between data sets (correlation between i, j > 0.79), completeness of 95.58%, and mean I/Sigma of 2.13. The final structure (PDB code: 9DKZ) was validated using the Comprehensive Validation Module (X-ray/Neutron)^70^ in Phenix including MolProbity^71^, real-space correlation, and atomic properties. The statistics associated with the structure are listed in Table 1. ChimeraX^72^ was used for analysis including electrostatic potential^73^ and molecular lipophilicity potential^74^ around macromolecules, and the generation of figures for visualization.

### Statistics

Statistical analysis for analysis in **Fig. 2d, e, g** was performed using Prism 10 (v. 10.1.0, GraphPad, USA). Normality of the distribution was first performed using Shapiro-Wilk test (*α* = 0.05). A *t* test (two tailed, *P* < 0.05, **Fig. 2d, g**) or one-way ANOVA (F = 22.38, DFn = 2, DFd = 59, *P* < 0.0001) was applied for normally distributed data (**Fig. 2e**).

### Data Availability

The MicroED structure of the eosinophil granular major basic protein-1 (gMBP-1) has been deposited to wwwPDB database under PDB accession code: 9DKZ.

## Author Contributions

E.R.W., J.E.Y., D.F.M., J.M.M. conceived the study. J.E.Y. and E.R.W. designed the study. J.E.Y. and J.M.M. prepared the samples and performed the experiments. J.E.Y. and C.A.B. processed the data. J.E.Y. and E.R.W. wrote the manuscript, with contributions from all authors. All authors read and approved the manuscript.

## Acknowledgement

This work was supported in part by the University of Wisconsin, Madison; the Department of Biochemistry at the University of Wisconsin-Madison; and grants U24 GM139168 to E.R.W., P01 HL088594 to Nizar Jarjour, and R01 AI125390 to D.F.M. and Joshua Coon from the National Institutes of Health. The work was in part supported by the grants DE-SC0023013 and DE-SC0018409 from US Department of Energy. J.M.M. was supported by the grant T32 HL07899 from the National Heart, Lung, and Blood Institute, National Institutes of Health. We are grateful for harvest and purification of donor-derived eosinophils by the eosinophil purification resource of the Department of Medicine, University of Wisconsin-Madison, and support from Sameer Mathur, M.D., Ph.D. and Paul Fichtinger. We are grateful for the use of facilities, instrumentation, and staff support at the Cryo-EM Research Center and the Midwest Center for Cryo-Electron Tomography in the Department of Biochemistry, University of Wisconsin, Madison. We are grateful for the computational resources supplied through the SBGrid. We are grateful for fruitful discussions with Francis Reyes, Ph.D. from ThermoFisher Scientific, and TEM and FIB/SEM instrumentation support from Micky Woods and Thomas Coomes from ThermoFisher Scientific, and the NIH-supported workshop “2022 MicroED Imaging Center Workshop at UCLA”.

## Extended figures and legends

**Extended Data Fig. 1.**
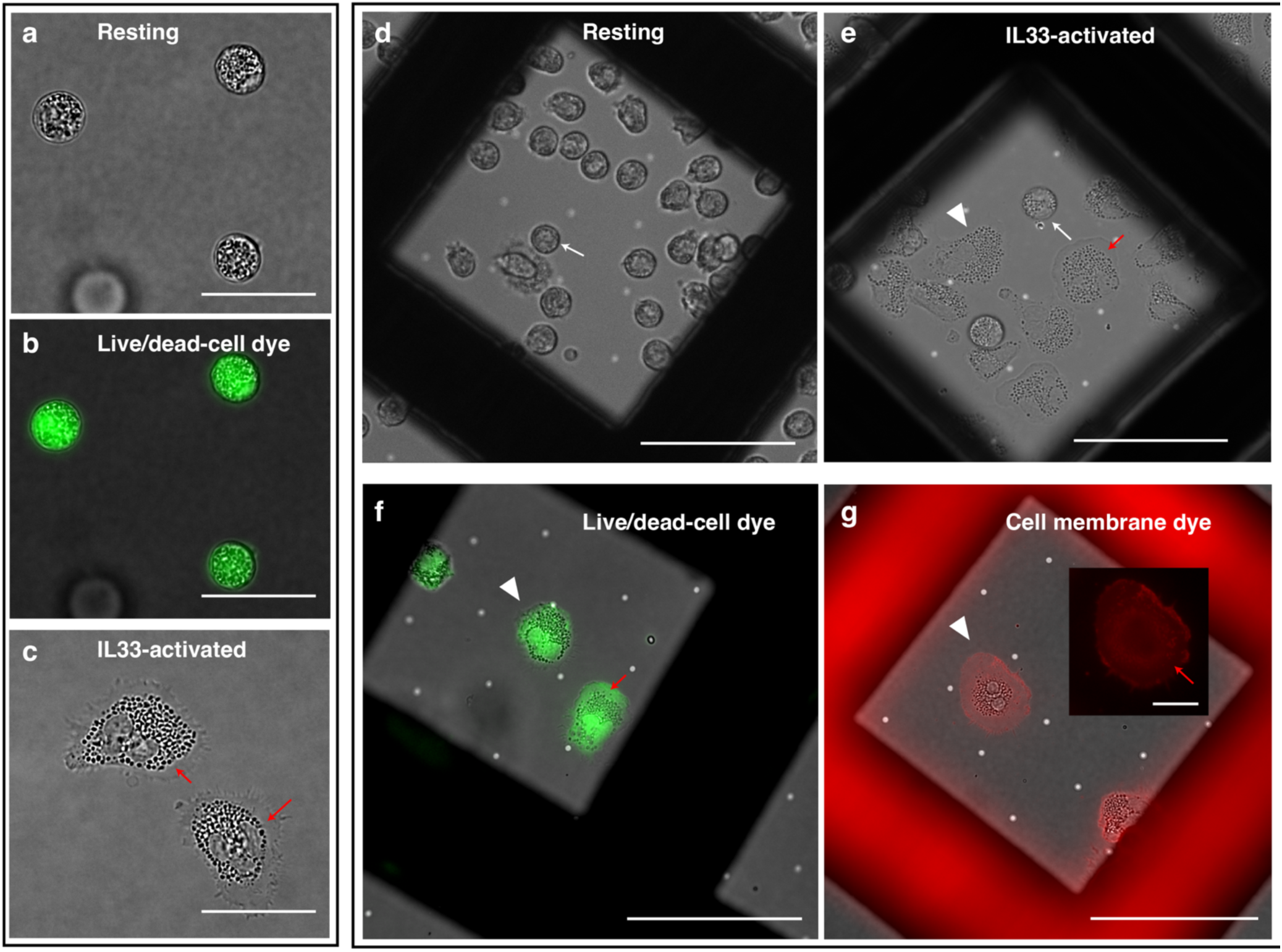
TEM grid preparation of resting and IL33-activated human eosinophils. **a-c,** Resting (**a-b**) and IL33-activated (**c**) donor-derived eosinophils exhibit the typical cellular morphology of rounded spheres (**a-b**) or flattened pancakes with extended cell edges (**c, e,** red arrow) on glass coverslips, respectively. Gentle adherence of eosinophils using fibrinogen onto the carbon-film of gold TEM grids minimized unwanted activation and preserved the resting states of the cells (**d**). Subsequent on-the-grid (fibrinogen-coated) IL33-treatment of eosinophils induced cell activation. Cells exhibited flattened pancake-like cell bodies packed with secretory granules (SGr) and flattened, thin extensions noted by red arrows (**c, e-g**). Cells were stained for either viability (green, Calcein live-cell dye, **b, f**) or cytoplasmic membrane continuity (**g**) to confirm that the inoculation of cells to EM grids and IL33-activation did not negatively impact cell health. Scale bars = 20 µm in **a-c**, 50 µm in **d-g**, 10 µm in the inset of **g**. White arrows, white arrowheads, and red arrows indicates resting eosinophils, IL33-activated eosinophil cells, and flattened extension from the IL33-activated eosinophil cell body, respectively.

**Extended Data Fig. 2.**
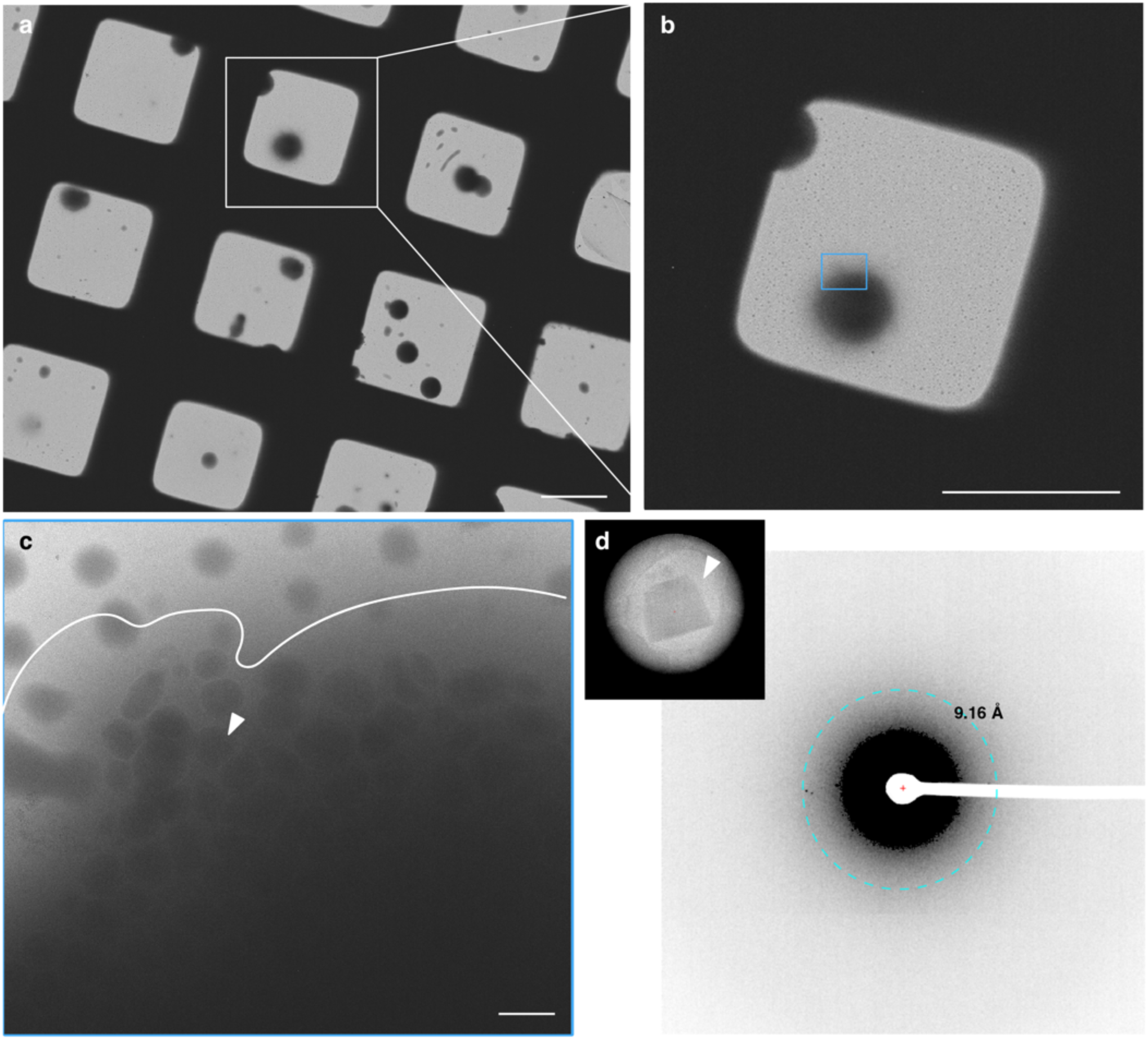
Low dose cryo-EM images of resting human eosinophils. **a-b,** A grid overview image (**a**) and an enlarged square view image at low magnifications (**b**) of resting donor-derived eosinophils distributed across the TEM grid after plunge freezing. The eosinophils were spherically shaped. Low dose 2D imaging at an intermediate magnification (EFTEM, 10 eV, 6500x, 38.52 Å/pixel) confirmed that the eosinophils were cryo-preserved with intact plasma membranes (**c**, delineated in white) and illustrated a denser cell interior associated with the resting state morphology. Weak diffraction (**d**, white arrowheads in **c** and **d**) from the intragranular nanocrystals in the resting eosinophils implied that the sample thickness was not suitable for direct electron diffraction studies. Scale bars = 50 µm in **a, b**, 10 µm in **c**.

**Extended Data Fig. 3.**
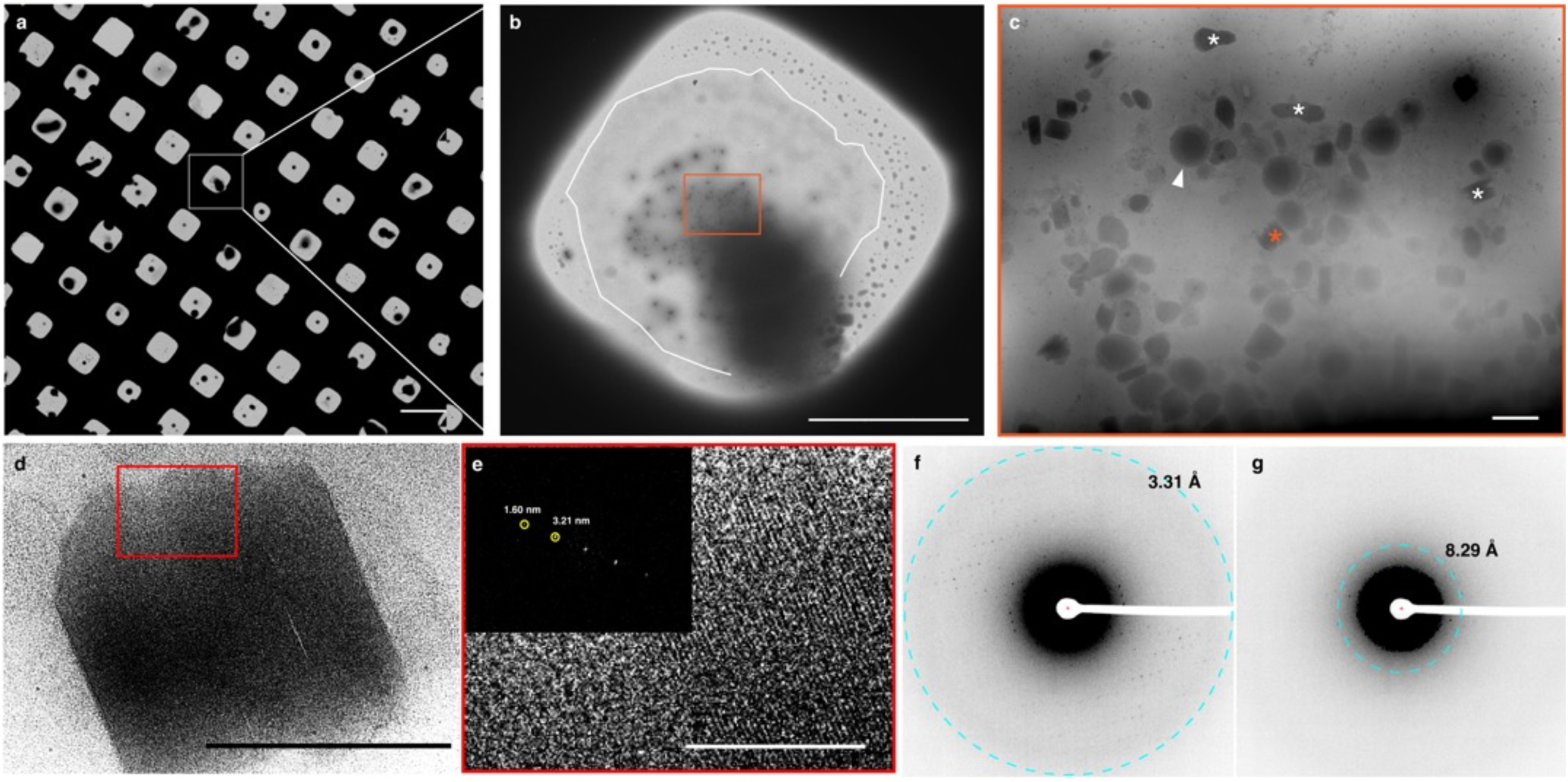
Low dose cryo-EM images of free crystalline cores in IL33-activated eosinophils. **a,** A grid square cryo-EM view of IL33-activated eosinophils showed that ∼60 % of the eosinophils were activated. A prototypical IL33-activated eosinophil was selected for the enlarged square view image (**b**). **b-c,** Both intact secretory granules (white arrowhead) and free crystalline cores (asterisk) were observed in cells that were activated by IL33 for 1 hour. **d-e,** Free crystalline structures were seen in the extended thin cell protrusions of activated eosinophils (thin membrane boundary delineated with a white line in **b**). 2D Fast Fourier Transfer (FFT) analysis of the low dose cryo-EM image of a representative free crystalline core (orange asterisk in (**c**)) showed clear lattice packing (**e**). Comparison of diffraction data collected at an accumulated dose of 0.05 e/Å^2^ (**f**) and 0.25 e/Å^2^ (**g**) of a representative free crystalline core. Scale bars = 100 µm in **a**, 50 µm in **b**, 1 µm in **c**, 500 nm in **d, e**.

**Extended Data Fig. 4.**
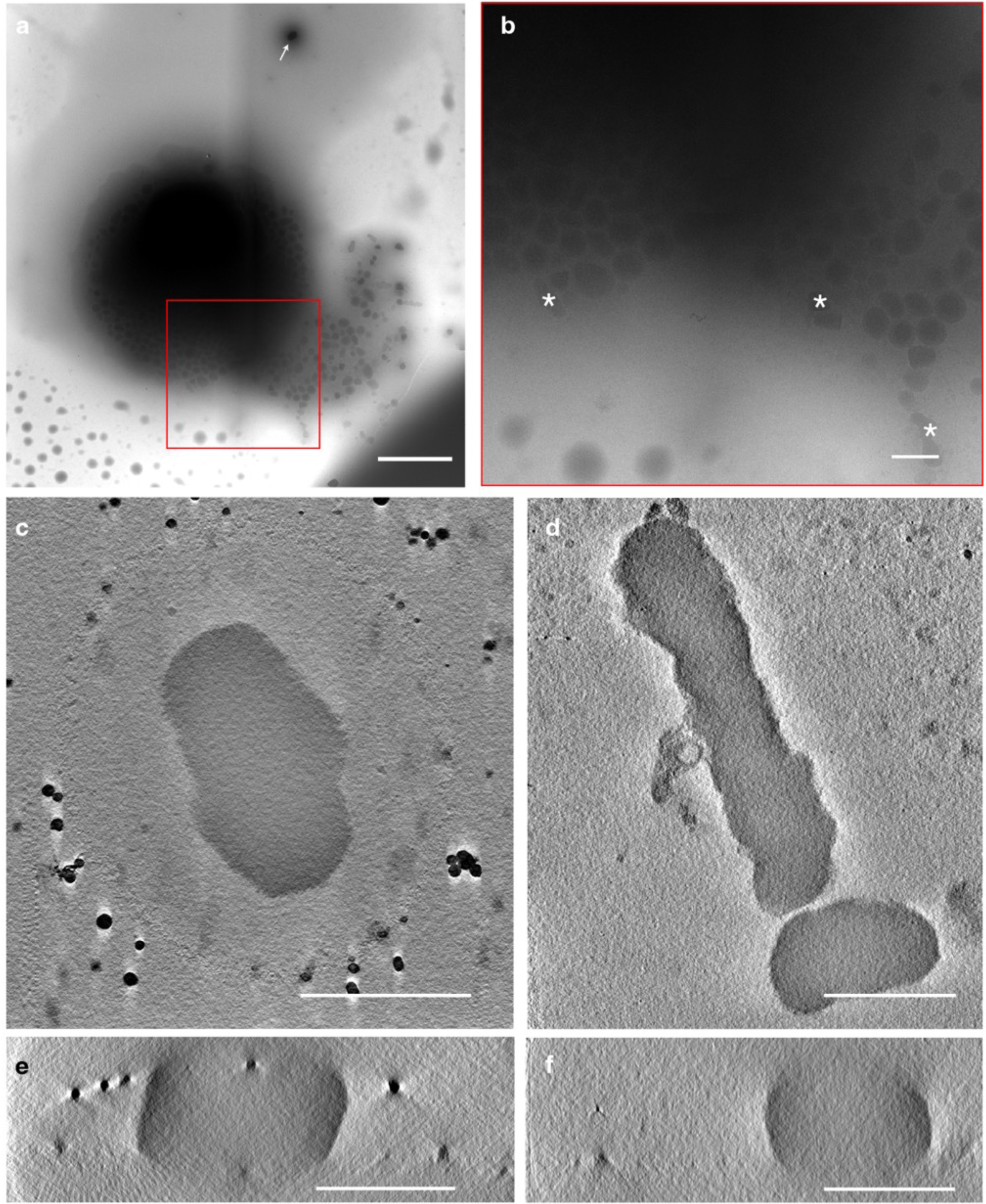
Cryo-ET of membrane-less free crystalline core from IL33-activated eosinophils. **a,** A low dose 2D montage view of an IL33-activated eosinophil with an extended zone of a barely discernable cell edge. **b,** Enlarged view (red boxed region in **a**) of free crystalline cores (white asterisks) in the extended edge zone. Nearby free crystals yet unassociated with the cell periphery were noted by white arrows. **c-f,** Tomographic slice views of free crystalline cores in XY (c, d) and XZ (e, f) demonstrate the membrane-less feature of these crystals and their size in 3D. Scale bars = 5 µm in **a**, 1 µm in **b-d**, 500 nm in **e, f**.

**Extended Data Fig. 5.**
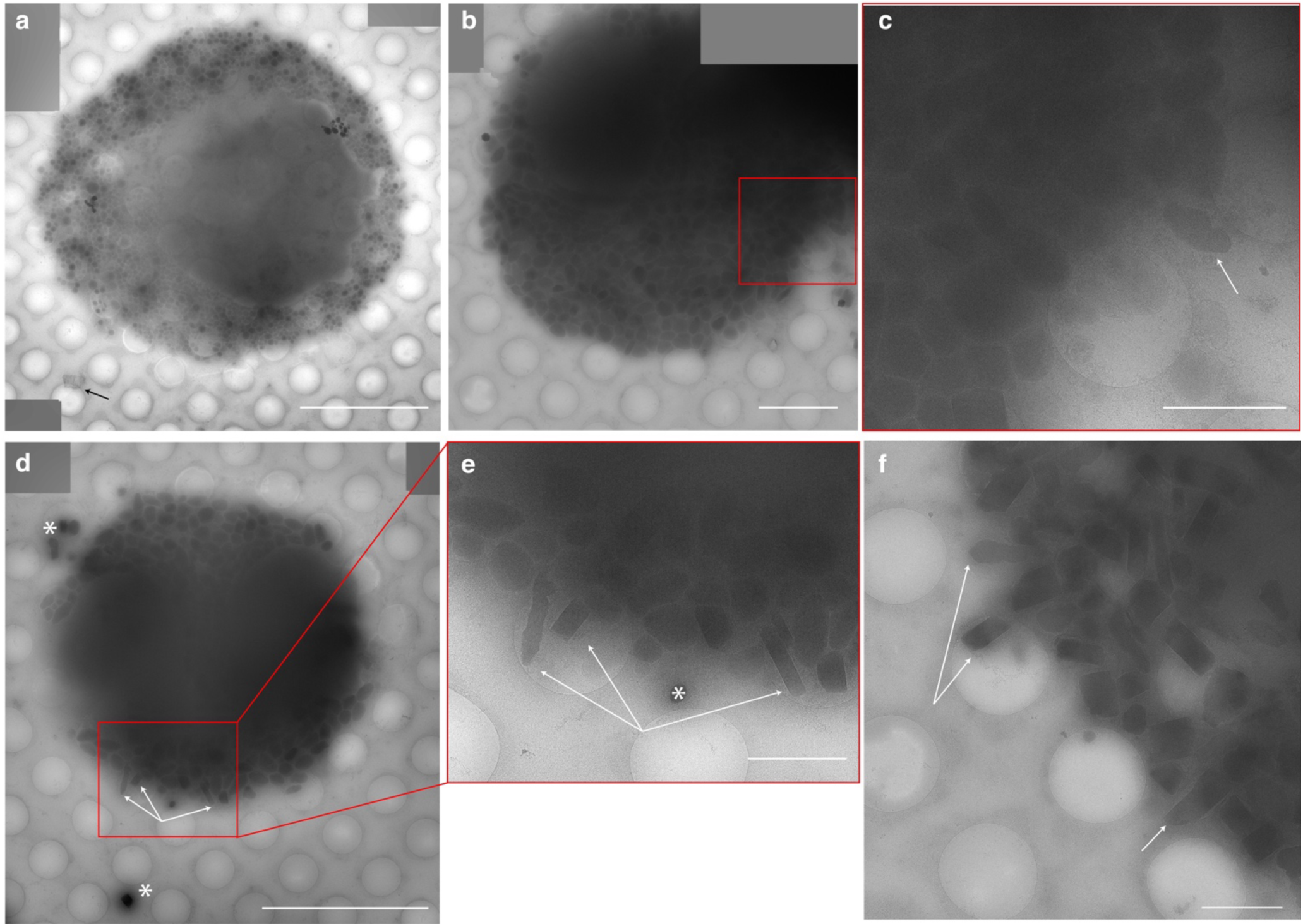
Cryo-EM characterization of IL33-activated eosinophils. **a**, IL33-activated eosinophil assumed a flattened overall shape with secretory granules and vesicular structures densely packed in the cytoplasm. Here, the activated eosinophils did not have clear thin cell extensions observed in Extended Data Fig. 1 and 3. Carbon foil debris were seen occasionally (black arrow in **a**). **b-f,** Snapshots of crystalline cores of secretory granule being released from the cell (white arrows) into the extracellular space were captured during the process of content exocytosis. **c** and **e** are enlarged views of the red boxed region in **b** and **d**. Free crystals (white asterisk) in proximity to or nearby the IL33-activated cell body were seen. Scale bar = 10 µm in **a, b,** 2 µm in **c, e, f.**

**Extended Data Fig. 6.**
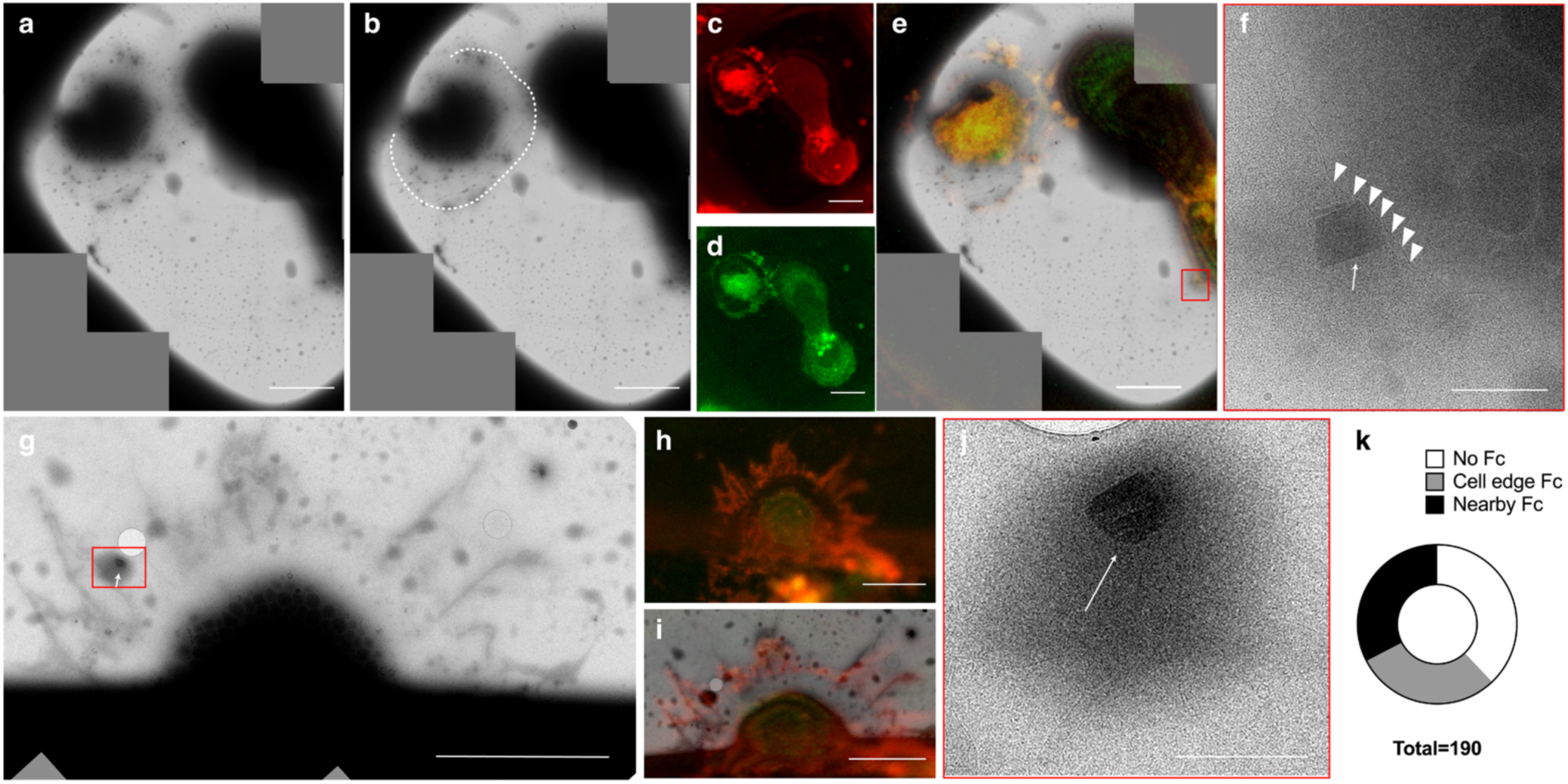
Extended thin cell protrusion in the IL33-activated eosinophils. Most (∼70%) of the activated eosinophils had an extended, protruding (**a, g**) membrane-enclosed zone, distinctly separated from the central cell body where secretory granules (SGr) were packed, as delineated in cryo-EM images (**b,** white dotted lines), and stained with low-toxicity cytoplasmic membrane dyes (red, **c, e, h, i**). As expected, the SGr had an innate green autofluorescence (**d, h, e**) due to the intragranular presence of flavins. Consistently, crystalline cores were seen attached to the cell membrane (**f,** white arrowheads) indicative of the release of SGr content. Free crystal cores were also observed in the extended membrane enclosed zone (**g,** white arrow). Statistical analysis showed that free crystals attached to the cell edge (**f**) including the ones in the thin extension zone (**g**), or in proximity of, yet not attached to the eosinophil (**j,** white arrow), were seen in roughly two-thirds (∼70%) of IL33-activated cells. Overlay of fluorescence light microscopy images and cryo-EM images (**c, i**) show the SGr packed cell body area and extruding membrane zone. Scalebars of 10 µm in **a, b, c, d, e, g, h, i,** 1 µm in **f** and **i.**

**Extended Data Fig. 7.**
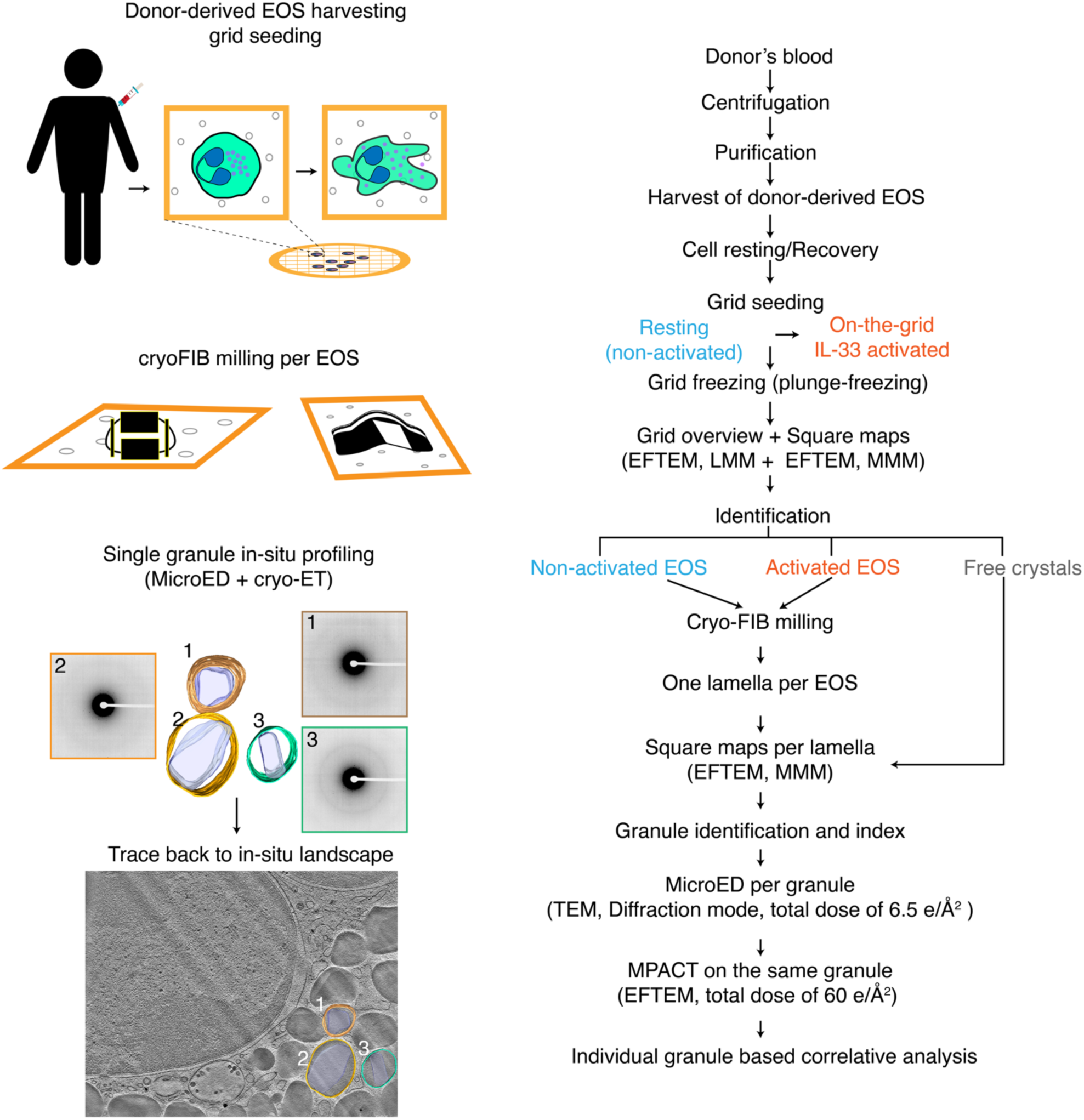
*In-situ* single granule profiling workflow. Donor-derived human eosinophils (EOS) were harvested and deposited on fibrinogen-coated TEM grids after a recovery resting phase. Prior to plunge freezing, half of the eosinophils were maintained in the resting state and the other half were activated with IL33 on the grid for 1 hour. The frozen grids were initially screened by low dose 2D cryo-EM to confirm and identify the status of activation based on cell morphology on a single cell level: resting (round) or IL33-activated (flattened). To study the structure of the cytoplasmic secretory granules (SGr), cryo-FIB milling was used to remove excess material, generating one thin lamella per cell. The milled grids were then examined under low dose EFTEM imaging (< 0.01 e/Å^2^) to locate milled cells and catalogue the visible SGr per cell. MicroED data was acquired on the catalogued SGr. Cryo-ET was subsequently collected on the same targets. Specifically, montage cryo-ET via MPACT was used to place the SGr in the native cellular landscape. As a result, each granule was correlatively profiled by both MicroED and cryo-ET. For the free, membrane-less crystalline cores present in flattened areas of IL33-activated eosinophils, correlative single crystal profiling was done directly without FIB-milling.

**Extended Data Fig. 8.**
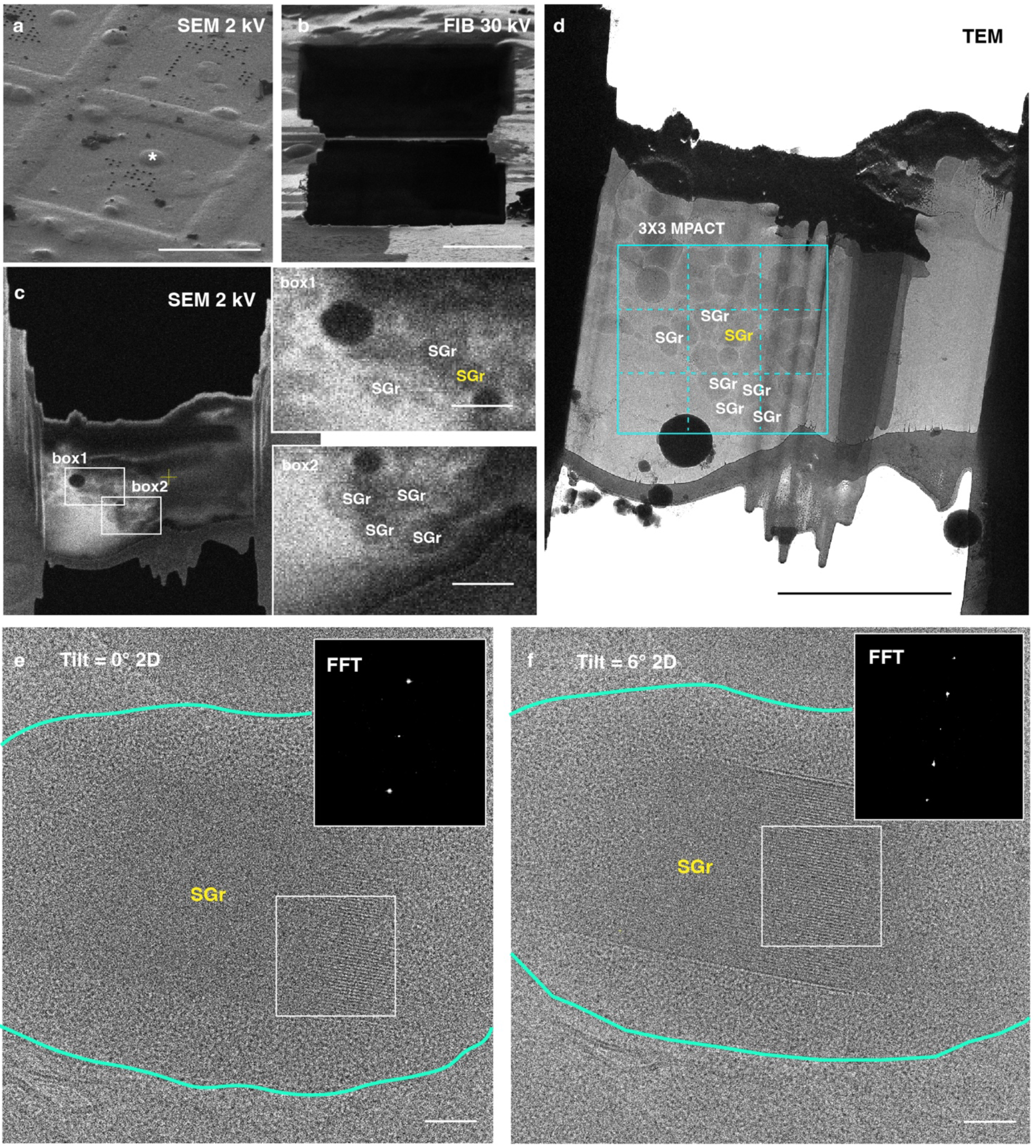
Cryogenic FIB fabrication and low-dose TEM imaging of human eosinophils in the resting state. Following initial low-dose cryo-TEM grid screening, individual grids with the resting eosinophils were loaded onto an Aquilos 2 cryo-FIB-SEM. **a**, Resting eosinophils, with rounded spherical morphologies were easily identified under cryo-SEM (stage tilt of 35°). The cells (white asterisk in **a**) were then FIB-milled using a gallium ion source. The FIB-milled lamellae were ∼220 nm thick (viewed under FIB, **b**). **c**, cryo-SEM imaging of the final 200-nm lamella revealed clear electron-dense vesicles that resembled secretory granules (SGr). Box1 and Box2: enlarged views of the boxed regions on the left. **d**, Low dose cryo-TEM of the same lamella and labeled SGr in the correlated cryo-SEM view (**c**). **d,** Acquisition of a montage tilt series via MPACT (3×3, solid cyan box, dashed lines indicative of individual tile frames). **e-f,** Enlarged cryo-EM views of the highlighted SGr and its nanocrystal core (yellow in **c, d**) at the tilt angle of 0° (stage tilt of 9°) and 6° (stage tilt of 15°), with corresponding Fast Fourier Transform (FFT) analysis of the white boxed region. The peaks correspond to 2.95 and 6.28 nm^-1^. Scale bars of 50 µm in **a**, 5 µm in **b, d**, 10 µm in **c**, left, and 1 µm in enlarged boxed views 1 and 2 **c**, right, 50 nm in **e-f**.

**Extended Data Fig. 9.**
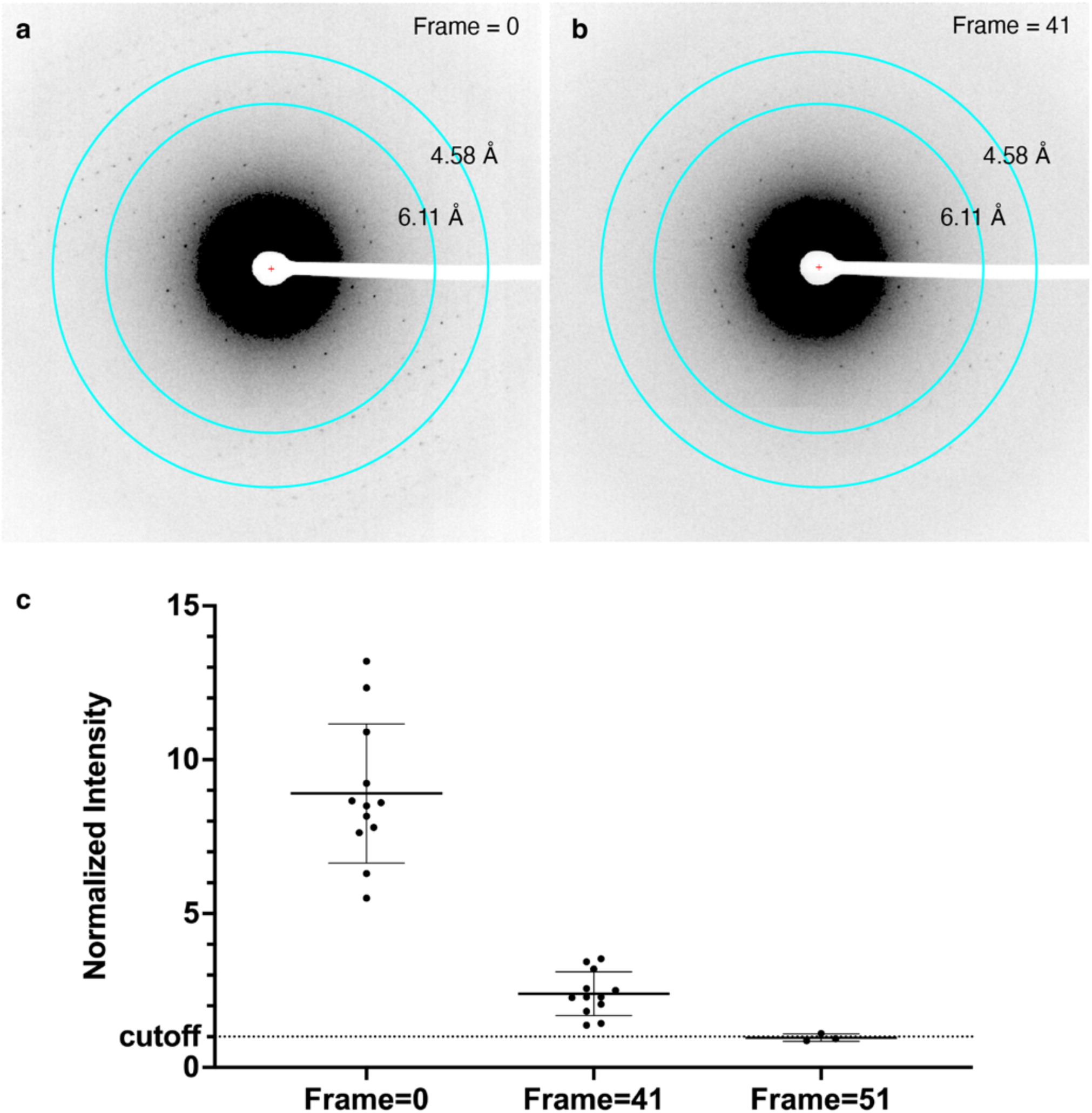
Critical electron dose determination for *in-situ* human major basic protein-1 from native secretory granule. Following cryo-FIB milling, an intact secretory granule with a clear nanocrystal core from a resting eosinophil was identified on the lamella under TEM. A single diffraction exposure of the selected granule was taken at the tilt angle of 0° (stage pre-tilt of 9°) with a dose of 0.15 e/Å^2^ (Frame = 0, **a**), followed by a continuous stage tilting MicroED acquisition to cover a range of 40° (N_frames_ = 40 frames) or 50° (N_frames_ = 50 frames) using the same dose (0.15 e/Å^2^) per exposure. After tilting, a final diffraction exposure of the same granule was collected at the tilt angle of 0° with a dose of 0.15 e/Å^2^ (Frame = 42, **b**). **c,** The total accumulated dose was ∼6.3 e/Å^2^ (N_frames_ = 40), or ∼7.8 e/Å^2^ (N_frames_ = 50). **c**, Normalized intensities (I_signal_/I_average_) of three diffraction spots in the resolution range of 4 to 7 Å were plotted at the initial exposure (Frame =0) and final exposure (Frame 41 or 51). The critical dose cutoff is the Normalized Intensity of 1 (dashed line), when a decrease in diffraction intensity becomes obvious. The determination of accumulated dose was repeated 4 times (N_frames_ = 40) on different intragranular nanocrystals.

**Extended Data Fig. 10.**
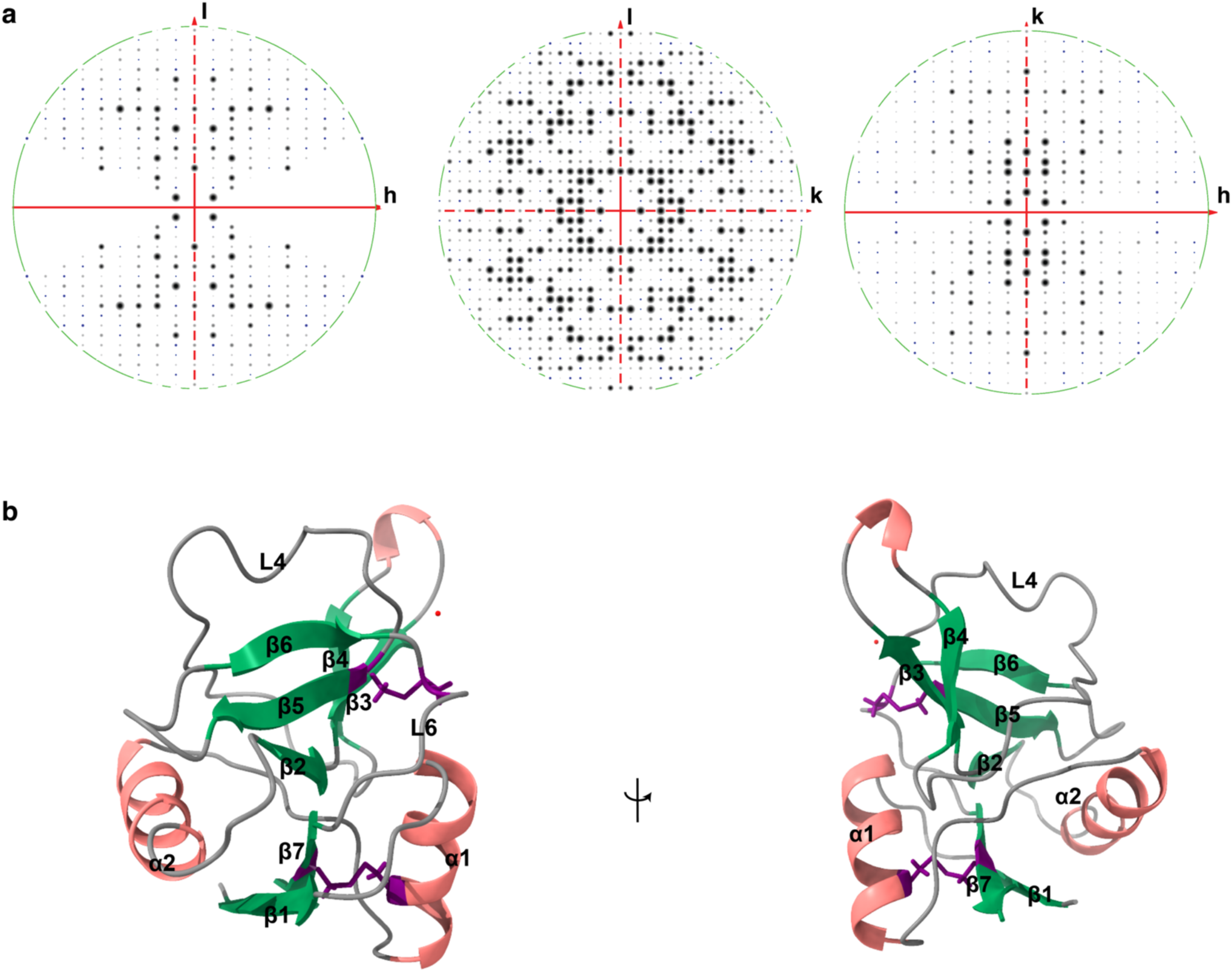
The structure of human eosinophil granular major basic protein-1 (gMBP-1). **a,** Completeness of the final merged map (n = 6) with the reflections viewed from the zones of *h*0*l*, 0*kl*, and *hk*0 showed the resolution cutoff (green circle) of 3.2 Å. **b,** The structure of gMBP-1 (PDB code: 9DKZ) in a ribbon model representation is comprised of two main α-helices (α1 and α2) and seven *β* strands (*β*1 – *β*7) connected by six loops (L1 – L6). Two disulfide bonds (Cys125-220 and Cys197-212), highlighted in purple, stabilized the overall topology of CTL domain. One water molecule (red dot) was modeled in with confidence at the current resolution cutoff.

**Extended Data Fig. 11.**
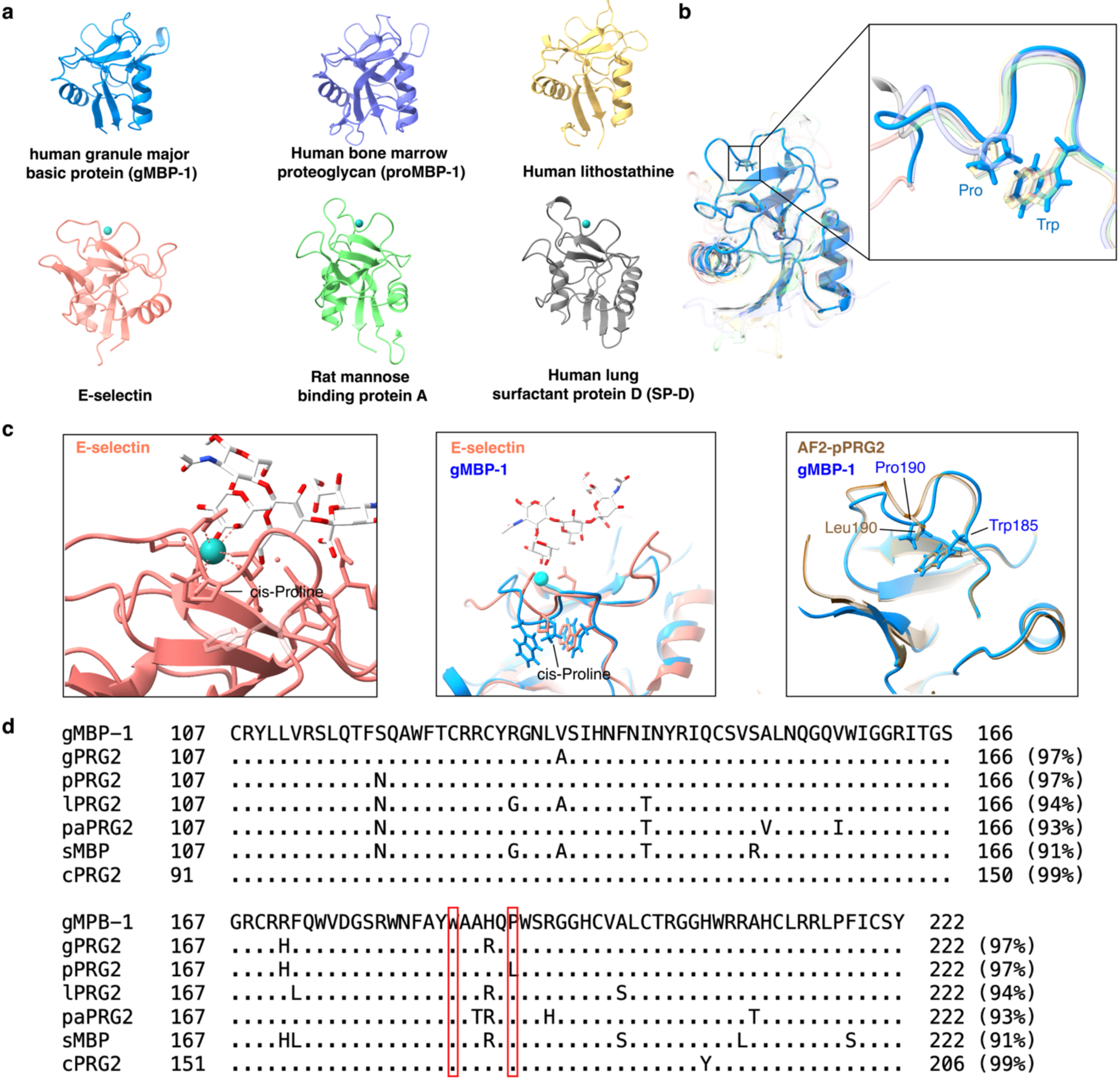
Comparison of *in-situ* human eosinophil granular major basic protein-1 (gMBP-1) with other C-type lectin (CTL) domain-containing proteins and homologues. **a-b**, Structures of CTL domain-containing proteins including, gMPB-1 (PDB: 9DKZ, blue), human bone marrow proteoglycan (proMBP-1/PRG2, PDB:8HGG, purple), human lithostathine (PDB: 1lit, yellow), human E-selectin (PDB: 1g1t, salmon), rat mannose binding protein A (PDB: 1kwt, green), and human lung surfactant protein D (SP-D, PDB: 1b08, grey) showed a similar overall topology (**a**) and calcium-mediated carbohydrate binding region (**b**) as observed in gMPB-1 (blue). Calcium was placed as a turquoise sphere in the binding pocket (**a** and **c**). **b,** An enlarged view of the binding pocket indicated the structural conservation of proline (Pro) and tryptophan (Trp) with overlayed gMBP-1 and other CTL domain containing proteins. **c,** Close inspection of the calcium binding region of E-selectin (c) indicates the role of Pro81 in creation of the pocket (left) for substrate accommodation. Pro190 in gMBP-1 contributed to the granular crystal packing through favored proline-aromatic interactions (middle). **d,** Sequence comparison of the CTL domain across bone marrow proteoglycan PRG2 homologues including Human (gMBP-1), Gorilla (gPRG2), Pan troglodytes (pPGR2), Hylobates lar (lPRG2), Pongo abelii (paPRG2), Symphalangus syndactylus (sPRG2), Chinese hamster (cPRG2). While pPRG2 has residue Leu190 the position of Pro190, the predicted pPRG2 (H2Q3N9) adopted a similar loop configuration as in gMBP-1 and other CTL domain proteins (**c,** right).

**Extended Data Fig. 12.**
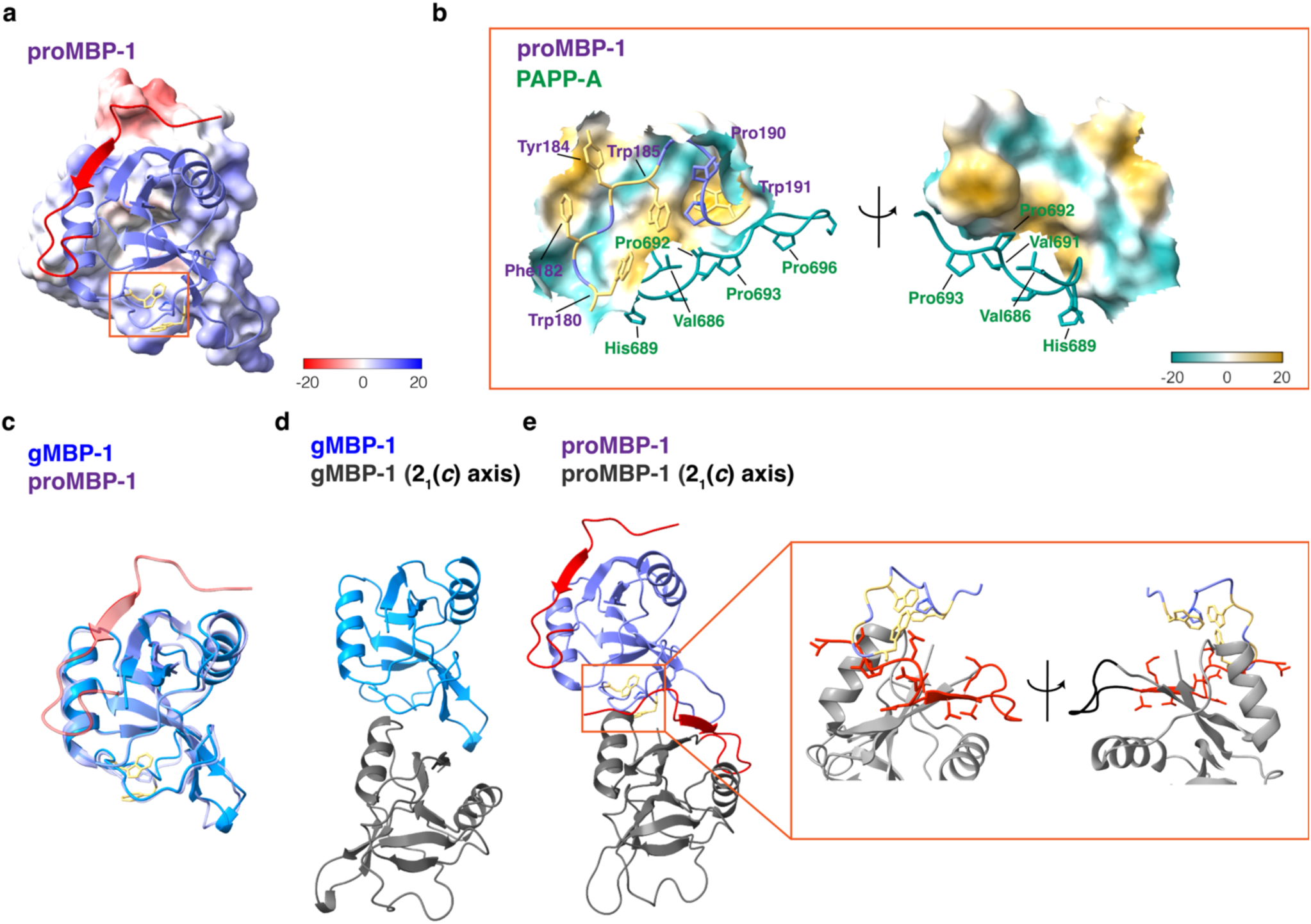
Structure of proMBP-1 and comparison with human eosinophil granular major basic protein-1 (gMBP-1). **a,** Ribbon illustration of proMBP-1 (PDB code: 8hgg, chain A) overlayed with the calculated electrostatic potential surface (red for negative potential through white to blue for most positive potential within a defined range of -20 to 20). Aromatic sidechains (yellow) of residues Trp185, Trp191 and residue Pro190 within the carbohydrate-binding loop region are highlighted in the boxed region (orange). **b,** Detailed view of the interactions in the proMBP-1 (purple) and PAPP-A (forest green) complex (PDB code: 8hgg, chain A and C). Multiple proline residues (Pro692, 693, 696, forest green) are seen with preferential contacts to the aromatic residues within the carbohydrate-binding loop region of proMBP-1 overlayed with the calculated molecular lipophilicity potential surface (dark cyan for the most hydrophilic, white and dark goldenrod for the most hydrophobic, on a scale range of -20 to 20). **c,** Superimposition of gMBP-1 (blue, PDB: 9DKZ) and proMBP-1 (purple) in a ribbon representation. The propiece (red) and the carbohydrate-binding loop region were highlighted (orange boxed). **d.** The crystal organization of two gMBP-1 monomers along the 21(*c*)-axis. **e.** The hypothetical crystal packing of two proMBP-1 monomers when adopting the same space group organization of gMBP-1 along the 21(*c*) axis. The presence of the propiece in proMBP-1 hinders the proper interactions between two monomers in the carbohydrate-binding loop region with detailed enlarged views of the red boxed region. The propiece residues from one monomer (grey ribbon representation) are colored in red while the critical aromatic residues Trp185, Trp191, Trp180 from the neighboring monomer along the 21(*c*) axis monomer are highlighted in yellow.

**Supplemental Table 1:** Donor characteristics and the number of grids prepared from purified Eosinophils. A total of 21 experiments were performed using Eosinophils from 16 unique donors (*repeat donors). Activation: IL33 or not treated (NT). Substrate: Fibrinogen (F). Dyes: Live/Dead dye (LD), CellMask (CM), CellBrite dye (CB), no dye (ND).

| Donor ID | Sex | Age | Asthma | Allergy | Activation | Substrate | Dyes | # of Grids |
| --- | --- | --- | --- | --- | --- | --- | --- | --- |
| BE0534 | M | 32 | Y | Y | IL33/NT | F | LD/CB/ND | 17 |
| BE0607 | F | 23 | Y | Y | IL33 | F | LD/CB/ND | 17 |
| BE0254 | M | 43 | Y | Y | IL33 | F | LD/CB/ND | 14 |
| BE0226* | F | 49 | Y | Y | IL33/NT | F | LD/CB/ND | 24 |
| BE0659* | M | 51 | Y | Y | IL33/NT | F | LD/CB/ND | 26 |
| BE0652* | F | 42 | Y | Y | IL33/NT | F | LD/CB/ND | 34 |
| BE0644 | M | 37 | N | Y | IL33 | F | LD/CB/ND | 12 |
| BE0628 | F | 53 | Y | Y | IL33 | F | LD/CM/ND | 12 |
| BE0656 | M | 46 | Y | Y | IL33 | F | LD/CM/ND | 12 |
| BE0660* | M | 33 | Y | Y | IL33 | F | LD/CM/ND | 17 |
| BE0622* | M | 45 | Y | Y | IL33/NT | F | LD/CB/ND | 27 |
| BE0545 | F | 54 | Y | Y | IL33/NT | F | LD/ND | 15 |
| BE0581 | M | 49 | N | Y | IL33 | F | LD/ND | 12 |
| BE0165 | M | 50 | Y | Y | IL33 | F | LD/ND | 8 |
| BE0281 | F | 50 | N | Y | IL33 | F | --- | 8 |
| BE0591 | M | 48 | Y | Y | IL33 | F | LD | 12 |

**Supplemental Table 2:**
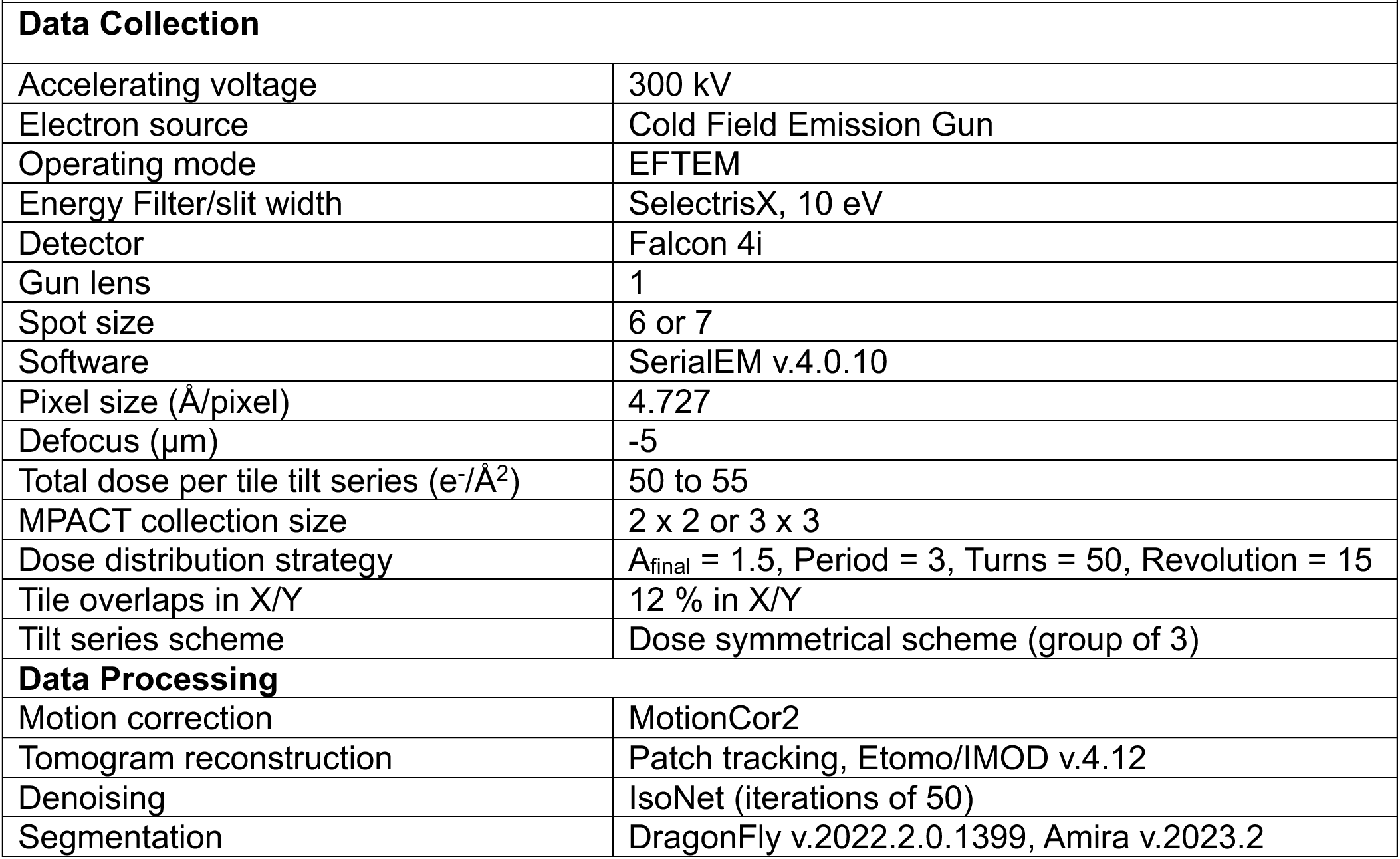
Montage Parallel Array Cryo-tomography (MPACT)

| **Supplemental Table 2. Montage Parallel Array Cryo-tomography (MPACT)** |  |
| --- | --- |
| **Data Collection** |  |
| Accelerating voltage | 300 kV |
| Electron source | Cold Field Emission Gun |
| Operating mode | EFTEM |
| Energy Filter/slit width | SelectrisX, 10 eV |
| Detector | Falcon 4i |
| Gun lens | 1 |
| Spot size | 6 or 7 |
| Software | SerialEM v.4.0.10 |
| Pixel size (Å/pixel) | 4.727 |
| Defocus (µm) | -5 |
| Total dose per tile tilt series (e<sup>-</sup>/Å<sup>2</sup>) | 50 to 55 |
| MPACT collection size | 2 x 2 or 3 x 3 |
| Dose distribution strategy | A<sub>final</sub> = 1.5, Period = 3, Turns = 50, Revolution = 15 |
| Tile overlaps in X/Y | 12 % in X/Y |
| Tilt series scheme | Dose symmetrical scheme (group of 3) |
| **Data Processing** |  |
| Motion correction | MotionCor2 |
| Tomogram reconstruction | Patch tracking, Etomo/IMOD v.4.12 |
| Denoising | IsoNet (iterations of 50) |
| Segmentation | DragonFly v.2022.2.0.1399, Amira v.2023.2 |

## Notes

### Competing Interest Statement

The authors have declared no competing interest.

